# Partitioning plant genetic and environmental drivers of above and belowground community assembly

**DOI:** 10.1101/173500

**Authors:** Matthew A. Barbour, Sonya Erlandson, Kabir Peay, Brendan Locke, Erik S. Jules, Gregory M. Crutsinger

## Abstract

1. Host-plant genetic variation affects the diversity and composition of associated above and belowground communities. Most evidence supporting this view is derived from studies within a single common garden, thereby constraining the range of biotic and abiotic environmental conditions that might directly or indirectly (via phenotypic plasticity) affect communities. If natural variability in the environment renders host-plant genetic effects on associated communities unimportant, then studying the community-level consequences of genetic variation may not be warranted.
2. We addressed this knowledge gap by planting a series of common gardens consisting of 10 different clones (genotypes) of the willow *Salix hookeriana* in a coastal dune ecosystem and manipulated natural variation in ant-aphid interactions (biotic) and wind exposure (abiotic) in two separate experiments. We then quantified the responses of associated species assemblages both above (foliar arthropods) and belowground (rhizosphere fungi and bacteria). In addition, we quantified plant phenotypic responses (plant growth, leaf quality, and root quality) to tease apart the effects of genetic variation, phenotypic plasticity, and direct environmental effects on associated communities.
3. In the ant-aphid experiment, we found that willow genotype explained more variation in foliar arthropod communities than aphid additions and proximity to aphid-tending ant mounds. However, aphid additions modified willow genetic effects on arthropod community composition by attracting other aphid species to certain willow genotypes. In the wind experiment, wind exposure explained more variation than willow genotype in structuring communities of foliar arthropods and rhizosphere bacteria. Still, willow genotype had strong effect sizes on several community properties of arthropods and fungi, indicating that host-plant genetic variation remains important. Across both experiments, genetic variation in plant traits was more important than phenotypic plasticity in structuring associated communities. The relative importance of genetic variation vs. direct environmental effects though depended on the type of environmental gradient (G > E_aphid_, but E_wind_ > G).
4. *Synthesis.* Our results suggest that host-plant genetic variation is an important driver of above and belowground biodiversity, despite natural variation in the biotic and abiotic environment.

## Introduction

Intraspecific genetic variation is a key driver of trait variation within host plants, which in turn can have cascading effects on associated species and entire communities of organisms (Antonovics 1992; Fritz & Price 1988; Lamit *et al.* 2016; Maddox & Root 1990). For example, genetic variation in the leaf chemistry of cottonwoods (Whitham *et al.* 2006) and in the plant architecture of coyote bush (Crutsinger *et al.* 2014) structures diverse assemblages of species, from foliar arthropods aboveground to soil microbes below. While community-level consequences of genetic variation (commonly referred to as ‘community genetics’, sensu Antonovics 1992) have been documented in a variety of host-plant taxa (Whitham *et al.* 2012), evidence comes primarily from common garden experiments. These controlled environments limit indirect effects of the biotic and abiotic environment on the expression of host plant traits (phenotypic plasticity, Gratani 2014) as well as direct environmental effects on the diversity and composition of species assemblages (Gaston 2000; MacArthur 1972).

To compare the community-level effects of host-plant genetic and environmental variation, researchers have taken two different approaches. A number of studies have planted common gardens in different environments across large spatial scales and measured community responses (e.g. Busby *et al.* 2014; Johnson & Agrawal 2005; Tack *et al.* 2010; Wagner *et al.* 2016). This approach has given important insight to processes such as local adaptation (Busby *et al.* 2014; Tack & Roslin 2010) and the scale-dependence of genotype-by-environment effects (Johnson & Agrawal 2005; Tack *et al.* 2010); however, the mechanisms underlying community assembly often remain poorly explored. In contrast, studies conducted at smaller spatial scales have manipulated specific biotic (e.g. Abdala-Roberts *et al.* 2012; Agrawal & Zandt 2003; Johnson 2008; Mooney & Agrawal 2008) and abiotic (e.g. Abdala-Roberts & Mooney 2012; Barrios-Garcia *et al.* 2016; Orians & Fritz 1996; Rossi & Stiling 1998) factors to gain detailed insight to community assembly. Although multiple biotic and abiotic factors can vary within a single ecosystem, most studies do not compare how multiple environmental forces and host-plant genetic variation structure associated communities.

Therefore, the importance of host-plant genetic variation in determining community assembly in naturally varying environments remains unclear for most systems (Crutsinger 2015; Hersch-Green *et al.* 2011; Tack *et al.* 2011).

Although host plants provide essential resources for a diverse array of taxa both above- and belowground, the majority of community genetics studies have focused on aboveground assemblages (Whitham *et al.* 2012). Studies that have simultaneously examined above- and belowground communities have found variable results, with host-plant genetic effects on aboveground communities being stronger (Bailey *et al.* 2009; Crutsinger *et al.* 2008; Wagner *et al.* 2016) or comparable and coupled (Crutsinger *et al.* 2014) with those belowground. Above- and belowground linkages can have important consequences for both plant fitness (Whitham *et al.* 2006) and terrestrial ecosystem processes (Wardle 2004). In addition, feedbacks between above- and belowground assemblages may depend strongly on the biotic and/or abiotic environment (Wardle 2004). Consequently, a rising challenge for community genetics is to understand the linkages between above- and belowground communities (Crutsinger *et al.* 2014; Lamit *et al.* 2015) and whether these linkages are modified by environmental variation.

Host-plant traits determine the quantity and quality of resources for the diverse organisms that colonize them; therefore, measuring functional trait responses of host-plant genotypes to different environments can give insight to mechanisms of community assembly. Phenotypic traits can vary in their plasticity (change in trait expression of a genotype in response to the environment: Scheiner 1993) and may even be plastic in response to one environmental gradient but not another (Garbutt & Bazzaz 1987; Scheiner 1993; Scheiner & Goodnight 1984). In addition, multiple plant traits can be important in structuring associated communities (Agrawal 2004, 2005; Agrawal & Fishbein 2006; Barbour *et al.* 2016, 2015). Simultaneous measurements of multiple functional traits and community-level patterns in genotype-by-environment studies can help distinguish the effects of genetic variation (proportion of variance in a trait explained by genotype: Lynch & Walsh 1998), phenotypic plasticity, and direct environmental effects on species assemblages. Yet, most genotype-by-environment studies focus on community responses and neglect to measure plant functional traits, preventing a mechanistic understanding of community assembly.

Here, we use common garden experiments to examine how host-plant genetic variation as well as the biotic and abiotic environment structure communities associated with the willow *Salix hookeriana* in a coastal dune ecosystem. Prior work in this system has shown that willow genotypes host distinct arthropod communities and that multiple plant traits determine community assembly (Barbour *et al.* 2016, 2015). Importantly, these traits varied substantially in their degree of heritability (plant growth, mean *H*^2^= 0.26; leaf quality, mean *H*^2^ = 0.72), suggesting that the environment may influence them in different ways. We sought to answer the following questions: (1) How do willow genotype and the environment affect plant functional traits and other environmental factors? (2) How do willow genotype and the environment affect the diversity and composition of above and belowground communities? (3) What is the relative importance of genetic variation, phenotypic plasticity, and direct environmental effects in structuring communities?

## Materials and Methods

### Study site

We conducted this research at Lanphere Dunes (40°53’29.85”N, 124°8’49.06”W), a restored coastal dune ecosystem managed by US Fish and Wildlife service in Humboldt County, California, USA. While most dunes on the Pacific coast of North America have become degraded due to a combination of development, use of off-road vehicles, and the invasion of non-native plant species, Lanphere Dunes has been restored and managed to be one of the last remaining areas of native dune left in California (Pickart 2013). Coastal willow (*Salix hookeriana* ex Barratt ex Hooker) naturally occurs in nearshore dune swales — seasonal freshwater wetlands that form in depressions between dune ridges (Pickart & Barbour 2007). Aside from coastal willow (hereafter willow), the dominant vegetation in these swales consists of beach pine (*Pinus contorta* ssp. *contorta*) and slough sedge (*Carex obnupta*).

During preliminary surveys, we qualitatively identified two important sources of environmental variation for willows in the dunes – one biotic (the presence of ant-aphid mutualisms) and one abiotic (wind exposure). We observed that the aphid *Aphis farinosa* was an abundant herbivore at Lanphere Dunes. *Aphis farinosa* is usually found at the tips of new shoot growth where it feeds on willow phloem. As with many other aphid species, *A. farinosa* excretes carbohydrate-rich honeydew while feeding, which attracts ants that tend the aphids and feed on the honeydew. This ant-aphid interaction is often mutualistic, because the ants will defend aphids from predatory arthropods and also eat other herbivores that may be competing with the aphids (Floate & Whitham 1994; Mooney & Agrawal 2008). The ant species we observed most frequently tending *A. farinosa* was the western thatching ant, *Formica obscuripes*. Western thatching ants create distinct dome-shaped mounds from nearby plant-material and are known to reduce herbivory from leaf chewing arthropods on *S. hookeriana* at our study site (Crutsinger & Sanders 2005), presumably by deterring ovipositing females or predating young larva. This work suggests that the presence of aphids and the proximity to ant mounds could influence associated communities through three non-mutually exclusive mechanisms: (i) increased abundance of aphid-tending ants, which could deter other arthropods; (ii) attraction of predators or deterrence of other herbivores, by aphids; (iii) alteration of plant-growth or leaf quality traits by aphids.

Wind exposure in coastal dunes can be an important environmental force in shaping plant growth traits and structuring communities associated with host plants (Crutsinger *et al.* 2014, 2010; Miller & Weis 1999). Preliminary surveys revealed that willows growing in wind-exposed habitats often exhibit reduced growth, especially at their leading edge, appearing to be “swept back” by the wind. We hypothesized that wind exposure may influence associated communities through three non-mutually exclusive mechanisms: (i) reduced plant size due to wind pruning; (ii) altered soil characteristics due to increased evaporation; and (iii) direct inhibition of ovipositing female arthropods.

### Experimental design

Prior to bud break in February 2012, we took shoot cuttings (40 cm length and ∼0.5 cm diameter) from one to two replicates of 10 different willow genotypes from a pool of 26 locally collected willow genotypes planted in a large common garden experiment in the same region as the current study site (∼24 km down the coastline). Details about the establishment of this common garden and genotyping are given in Barbour *et al.* (2015). These 10 genotypes displayed substantial variation in both plant-growth and leaf-quality traits (Barbour *et al.* 2015). Shoot cuttings were soaked in water overnight and then planted in a mixture of 80% perlite, 20% peat moss (dolomite lime added to balance pH) inside “Cone-tainers” (Stuewe & Sons, Inc.). We grew cuttings under ambient weather conditions outside the greenhouse at Humboldt State University until we transplanted willows into multiple common gardens at Lanphere Dunes.

### Ant-aphid experiment

To examine how the presence of aphids, proximity to ant mounds, and willow genotype affected associated communities, we established common gardens around 5 different ant mounds (treated as blocks) in late May 2012. Within each block, we randomly planted 20 cuttings (2 replicates of each of 10 genotypes) with 0.5 m spacing in plots that were at a distance of 1, 6, and 12 meters from the edge of the ant mound, for a total of 60 cuttings per ant mound (300 cuttings for entire experiment). Within each plot, we randomly assigned the aphid treatment (aphid presence vs. absence) to one of the two replicates for each genotype. On May 22, we collected aphids (*Aphis farinosa*) from a single willow patch at Lanphere Dunes and placed 5 adult apterate aphids on the tips of willow cuttings in the aphid treatment using a moist paintbrush. We bagged aphids onto the apical shoots of cuttings using organza bags to promote aphid establishment on plants. Similarly, we placed organza bags on all control plants. On May 27, we checked aphid treatments to ensure there were 5 adult aphids and removed bags from all cuttings. If necessary, we added aphids to these treatments until there were 5 adults and we removed any aphid nymphs that were produced since initial establishment. We checked plants for aphids on June 6, June 13, June 24, July 4, July 14, and July 20, 2012. If plants in the aphid treatment had less than 5 aphids, we noted their abundance and added aphids until there were at least 5 individuals. The ant-aphid experiment was restricted to the summer of 2012, because in the summer of 2013 there was high willow mortality induced by drought and *A. farinosa* was too low in abundance on naturally occurring willows to allow us to repeat the experiment.

### Wind experiment

To examine how wind exposure and willow genotype affected associated communities, we planted 200 willow cuttings in a split-plot experimental design in late May of 2012. At each of 10 different naturally occurring willow stands (treated as blocks), we established an ‘exposed’ and an ‘unexposed’ common garden with exposed gardens on the windward side of natural willow stands and unexposed plots on the leeward side. Each garden consisted of one replicate cutting of each of 10 genotypes randomly planted in 2 m by 0.5 m grid with 0.5 m spacing between plants. The center of exposed and unexposed gardens within each block were the same distance (2 m) from the edge of the willow stand to control for insect accessibility. To estimate the difference in wind conditions experienced by exposed vs. unexposed plants, we went out on a representative windy afternoon in September 2012. A nearby weather station (Arcata, CA) estimated wind speeds of 22 km/h during this period. We used a hand-held anemometer (Kestrel 1000) to measure wind speed at a height of 37 cm aboveground (approximate height of tallest plants in the garden in 2012) in each plot of our experiment. For each block, we randomly selected the order in which exposed and unexposed plots were measured and took maximum wind speed measurements over a 30 s period. We found that willows growing in wind-exposed plots experienced up to 3.7-fold higher wind speeds compared to unexposed plots (*F*_1,9_ = 187.32, *P* < 0.001), suggesting that the location of our plots were effective manipulations of wind exposure.

### How do willow genotype and the environment affect plant functional traits and other environmental factors?

#### Plant traits

Prior work in this study system demonstrated that variation in both plant growth and leaf quality traits affects the community assembly of herbivorous insects (Barbour *et al.* 2015). To quantify plant-growth traits, we measured plant height, the number of shoots produced, and average shoot length in late July of each year (end of growing season) for both experiments. We quantified plant height as the distance (mm) from the ground to the tip of the tallest shoot. We quantified average shoot length by measuring every shoot on each plant to the nearest millimeter and calculating the average shoot length for each plant. We also measured several traits that could affect leaf quality for herbivores, including water content, trichome density, specific leaf area (SLA), percent carbon (C) and nitrogen (N), and C:N. To measure these traits, we excised fully expanded and undamaged leaves from plants in late July of each year, stored leaf samples with a moist paper towel in separate plastic bags within a cooler and immediately brought them back to the laboratory. We then weighed leaves to obtain fresh mass (g), digitally scanned them to measure leaf area (*mm*^2^) using ImageJ (Abràmoff *et al.* 2004), and oven-dried them at 60 °C for 72 h to obtain dry weight (g)(Cornelissen *et al.* 2003). We calculated SLA as (*leaf area*) / (*dry mass*) (Cornelissen *et al.* 2003). We calculated leaf water content as the (*fresh mass* – *dry mass*)/(*dry mass*) (Cornelissen *et al.* 2003). To measure trichome density, we counted the number of trichomes along an 11 mm by 1 mm transect in the center of the leaf, halfway between the leaf edge and the mid-vein, under a dissecting scope. To measure percent C and N, we ground oven-dried leaves to a fine powder using a ball mill (Mixer/Mill 8000D, SPEX SamplePrep; Metuchen, NJ, USA). Subsamples of each material were then analyzed for percent C and N on an elemental analyzer (ECS 4010; Costech Analytical Technologies, Valencia, California, USA) using atropine (4.84% N and 70.56% C) as a reference standard. For root-associated communities, we hypothesized that variation in root C:N may affect community assembly. We measured root C and N by crushing a subsample of oven-dried roots with a razor blade and then analyzing for percent C and N on an elemental analyzer (Carlo-Erba NA 1500) using atropine (4.84% N and 70.56% C) as a reference standard.

Our prior work with *Salix hookeriana* has shown that it exhibits subtantial genetic variation in a suite of phenolic compounds that are known to influence the assembly of its herbivore community (Barbour *et al.* 2015); however, measuring phenotypic variation in these compounds across both experiments was beyond the scope of this study. Community genetic effects not explained by the plant traits we measured could be explained by heritable variation in these chemical compounds. To explore this, we used chemical data collected from the same genotypes growing in the large common garden (Barbour *et al.* 2015) where we obtained willow cuttings for the current experiments. While these data do not incorporate biotic and abiotic effects on willow phenolic chemistry, our prior work found that these compounds were highly heritable (mean *H*^2^ = 0.77) and therefore unlikely to be strongly influenced by environmental variation.

#### Soil characteristics

Soil nutrients, total organic matter, and moisture may all influence plant traits and the assembly of fungal and bacterial communities on plant roots (Erlandson *et al.* 2015). Moreover, we expected that wind exposure would affect these soil characteristics (Lortie & Cushman 2007); therefore, we measured soil nutrients, percent organic matter, and moisture within each plot of the wind experiment (one exposed and one unexposed plot per block). To estimate soil nutrient uptake by willows, we installed Plant Root Simulator (PRS) Probes (Western Ag Innovations, Saskatchewan, Canada) at three randomly selected locations within each plot for 11 days in September 2012. PRS Probes estimate nutrient supply rates to roots by continuously adsorbing charged ionic elements over the burial period. For our study, we estimated potential root uptake of 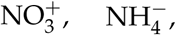 Ca^2+^, Mg^2+^, K^+^, 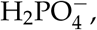 Fe^3+^, Mn^2+^, Cu^2+^, Zn^2+^, 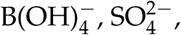 Pb^2+^, Al^3+^, Cd^2+^. From this nutrient data, we calculated total N as 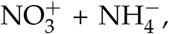 and then used principal components analysis to condense all nutrient variation into a single axis (nutrients PC1) that explained 34% of the variation. Nutrients PC1 described the negative correlation between nitrogen compounds 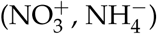 and the rest of the ionic elements, with positive values indicating high supply rates of all ionic elements except for the nitrogen compounds. To measure percent organic matter content (%OM), we used a trowel to collect soil (depth = 0 – 15 cm) adjacent to the randomly positioned PRS probes in September 2012. Soils were transported back to the lab in plastic bags, sieved into fragments less than 2 mm, randomly subsampled using a soil splitter, and dried at 105 °C for 72 hours. We then weighed a subsample of the oven dried soil into an oven dried crucible and placed the crucible and soil into a furnace to be combusted at 375 °C for 16 hours. We then weighed the mass (g) of combusted samples, placed them in a desiccator for 20 minutes, and weighed them again. We calculated percent organic matter as %*OM* = (*oven dry mass* – *combusted dry mass*) / (*oven dry mass*) × 100. To measure soil moisture (volumetric water content, m^3^/m^3^), we used a 5TE soil sensor coupled to an EM50 Digital/Analog Data Logger (Decagon Devices, Pullman, Washington, USA). In September 2012, while PRS probes were in the ground, we measured soil moisture at a depth of 5 cm in three random locations within each plot on three different days between 1100 – 1500 hours. We repeated this same sampling scheme in early July 2013. Plot levels measurements of soil moisture were highly correlated between years (Pearson’s *r* = 0.93, *t*18 = 10.91, *P* < 0.001), so we averaged these soil moisture estimates to determine a single soil moisture value per plot.

#### Analyzing plant trait and environmental responses

To analyze how willow genotype, the environment, and their interaction, influenced willow phenotypes, we used separate generalized linear mixed-effect models (GLMMs) (Bolker *et al.* 2009). For the ant-aphid experiment, we specified block (ant mound) and plots nested within block (the 3 different distances from ant mound) as random effects. We specified willow genotype, aphid treatment, distance from ant mound, and their 3-way interaction as fixed effects in the model. For the wind experiment, we specified block (willow stand) and plots nested within block (the 2 wind exposure treatments) as random effects. We specified willow genotype, wind treatment, sampling year, and their 3-way interaction as fixed effects in the model. We lacked multiple years of data on leaf trichome density (2012 only), SLA (2013 only), leaf C:N (2013 only), and root C:N (2013 only) for the wind experiment; therefore, we removed sampling year, and its interactions, from the fixed effects structures of these GLMMs. Plant mortality in each experiment resulted in unbalanced designs, so we used Type II sum-of-squares to test the significance of fixed effects. For continuous responses (e.g. plant height) we specified Gaussian error distributions in our models and tested the significance of fixed effects using *F*-tests with Kenward-Roger approximated degrees of freedom. For count responses (e.g. shoot count), we specified Poisson error distributions in our models and tested the significance of fixed effects using likelihood-ratio tests.

To examine the effect of wind exposure on soil characteristics (total N, nutrients PC1, %OM, and soil moisture), we used separate mixed effect models with wind treatment as a fixed effect and block (willow stand) as a random effect. Since all soil characteristics were continuous responses, we specified Gaussian error distributions in our models and tested the significance of fixed effects using *F*-tests with Kenward-Roger approximated degrees of freedom.

### How do willow genotype and the environment shape the diversity and composition of above and belowground communities?

#### Aboveground: foliar arthropods

We visually surveyed plants for arthropods to determine the abundances of different morphospecies. For the ant-aphid experiment, we surveyed arthropods on 5 different occasions between early June and late July 2012. For the wind experiment, we surveyed arthropods once at the end of July 2012 and then once a month in May, June, and July of 2013. So that individuals were not counted twice between sampling dates, we took the maximum abundance for each arthropod morphospecies from each plant across all sampling dates within each year. This approach provides a conservative estimate of the total number of individuals of each morphospecies that occurred on individual plants through the summer. Given the relatively low abundances of individual morphospecies, we grouped arthropods at the level of family for insects and order for all other arthropods prior to analyzing community composition (details in *Analyzing community responses* section below).

#### Belowground: rhizosphere fungi and bacteria

We dug up the willows from the wind experiment to sample fungal and bacterial communities associated with willow roots in late July of 2013. We were unable to sample belowground communities of plants in the ant-aphid experiment due to the high mortality of plants in 2013. To sample these belowground communities, we removed willows with the surrounding soil intact to preserve root systems, separated shoots and roots, then brushed soil off root systems and stored roots in separate plastic bags. Within 6 hours of excavation, root systems were stored at 4°C. To process roots, we gently rinsed them in tap water until free of visible soil. In order to randomly select roots for molecular analysis, second order roots were cut into 2 cm lengths, spread out on a grid, and then, using a random number generator, a total of 30 cm of root length was picked from numbered grid cells. These random root subsamples were flash frozen in liquid N, and kept at −80°C until DNA extraction. To increase efficiency of DNA extraction, roots were physically disrupted with 2 beads per 2 mL tube (3.0 mm Yttria stabilized Zirconia Grinding Media) for 30 seconds at 1500 strokes per minute (SPEX SamplePrep 200 geno/grinder). Total DNA was extracted from the root samples using MoBio PowerSoil 96 sample DNA extraction kits following the manufacture’s instructions.

To identify fungal and bacterial OTUs, we used custom Illumina-compatible barcode primer sets ITS1F/ITS2 (Smith & Peay 2014; White *et al.* 1990) and 515f/806r (Caporaso *et al.* 2012) to amplify via PCR the first internal transcribed spacer (ITS1) of the fungal nuclear ribosomal RNA operon and the V4 region of bacterial 16S ribosomal DNA from total root DNA extractions. Product quality was assessed by gel electrophoresis. PCR products were cleaned with house-made magnetic bead solution, quantified with a Qubit fluorometric kit, then sample libraries were pooled at a fungi:bacteria concentration ratio of 2:1. Pooled amplicon libraries were sequenced as single-index (the reverse barcode was uniquely indexed) 300 base pair reads at Stanford Functional Genomics Facility on one lane of an Illumina MiSeq. Quality control of reads consisted of these steps: trimming bases with quality score less than 20 phred; trimming sequenced adaptors; and removing reads with average error rates greater than 0.25 using UPARSE (Edgar 2013). Only high quality, paired forward and reverse reads were used for OTU clustering at 97% identity, then OTUs were checked for chimeras against the GOLD 16s rRNA database (Reddy *et al.* 2015) and UNITE fungal ITS database ver6_97_13.05.2014 (Kõljalg *et al.* 2005) with UPARSE. Taxonomy was assigned in QIIME (Caporaso *et al.* 2010) using the RDP Classifier for bacteria (Wang *et al.* 2007) and BLAST for fungi. We normalized datasets to account for differences in each sample’s library size (number of reads obtained for each sample). Finally, we discarded some OTUs and samples based on the following conditions: OTUs with no known taxonomy (any OTU that was not assigned to at least Kingdom Fungi, Bacteria or Archaea); root samples with fewer than 6000 fungal reads and 9000 bacterial reads; and mitochondrial and chloroplast OTUs.

#### Analyzing community responses

To determine how willow genotype, the environment, and their interaction, influenced richness, abundance, and rarefied richness of aboveground arthropods as well as rarefied richness of root fungi and bacteria, we used separate GLMMs with the same structure described for *Analyzing plant trait and environmental responses.* For the ant-aphid experiment, we omitted *A. farinosa* and *F. obscuripes* from our calculations of arthropod community properties because we expected our treatments to manipulate their abundances. For continuous responses (rarefied richness) we specified Gaussian error distributions in our models and tested the significance of fixed effects using *F*-tests with Kenward-Roger approximated degrees of freedom. For count responses (richness and arthropod abundances), we specified Poisson error distributions in our models and tested the significance of fixed effects using likelihood-ratio tests. If necessary, we accounted for overdispersion in these Poisson models by specifying an individual-level random effect.

To examine whether community composition depended on willow genotype, the environment, or their interaction, we applied a Hellinger transformation to our community data (square root of proportional abundance of species found on each willow; Legendre & Gallagher 2001) and conducted separate redundancy analyses (RDA, 1000 permutations on Euclidean distances) for the arthropod, fungal, and bacterial communities. A Hellinger transformation was appropriate because calculating Euclidean distances on raw abundance data can sometimes result in two communities that do not share the same species being more similar than two communities that do share the same species (Legendre & Gallagher 2001). We incorporated the same fixed effects structure as we used to analyze the univariate community responses for each experiment. To test the significance of each effect, we used Type II sum-of-squares and compared the observed community dissimilarities to the dissimilarities we would expect by random chance with a permutation test that controls for the blocked design of our experiment. To test the significance of treatments that varied at the plot-level (wind exposure and distance from ant mound), we first calculated the community’s centroid in multivariate space for each plot. We then included block as a covariate and ran the same permutation test as previously described. This ensured that our significance tests of treatments that varied at the plot-level were based on the appropriate residual degrees of freedom (wind exposure residual *df* = 9; distance from ant mound residual *df* = 4).

To compare the relative importance of willow genotype vs. different environmental factors, we fit reduced GLMMs with significant factors as random effects. We then used the variance components estimated for these random effects to calculate the percentage of variance (hereafter *σ*^2^) explained by willow genotype and specific environmental factors. This method has been proposed as appropriate for comparing the relative importance of plant genotype and the environment in genotype-by-environment studies (Hersch-Green *et al.* 2011; Johnson & Agrawal 2005). For models of community composition, we fit reduced RDAs with only significant factors and calculated the adjusted redundancy statistic for each significant factor, which has been shown to be an unbiased estimator of explained variance (Peres-Neto *et al.* 2006). This adjusted redundancy statistic is an analogue of adjusted *R*^2^ but for multivariate responses and we refer to it hereafter as 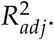

##### What is the relative importance of genetic variation, phenotypic plasticity, and direct environmental effects in structuring communities?

To tease apart the relative importance of these factors, we used piecewise structural equation models (SEMs, Lefcheck 2015). We modeled only potential mechanisms and community responses that were statistically significant in our prior analyses. For example, if we did not observe a G×E effect on a community response variable, then we did not model this interactive effect in the piecewise SEM. An advantage of piecewise SEMs is that they are flexible, allowing users to account for correlated structure (i.e. random effects) in their experimental design. However, as with any technique that relies on multiple regression, structural equation models can give misleading results if there is collinearity among predictor variables. To mitigate the effects of collinearity, we used principal components analysis (PCA) to condense aboveground willow phenotypes as well as soil properties into a small number of uncorrelated variables. For above-ground willow traits in the wind experiment, we analyzed separate PCAs for 2012 and 2013 since we did not always have data on the same traits in each year. At times, we lacked data for all traits on each plant or all soil properties measured in each plot. Therefore, we used a regularized iterative PCA algorithm to impute missing values (Josse *et al.* 2012). For each PCA, we retained principal components with eigenvalues greater than 1.

To calculate standardized coefficients (*β*) in our piecewise SEM, we scaled all predictor and response variables to mean = 0 and SD = 1 prior to analyzing them with GLMMs (error distribution = Gaussian). For willow genotype, we specified the average effect for the 10 genotypes as the reference level (i.e. deviation contrasts) and calculated the standard deviation of the coefficients to determine its standardized coefficient. We then multiplied standardized coefficients across a given pathway to calculate the strength of each mechanism we modelled. To evaluate the explanatory power of our separate GLMMs, we report marginal *R*^2^ (Nakagawa & Schielzeth 2012). Marginal *R*^2^ values do not adjust for the variance explained by our random effects; therefore, they give us a truer sense of the explanatory power of our models. To evaluate the fit of the full structural equation model, we used a test of directed separation (Shipley 2000). This test identifies missing paths in the model, calculates the P-value for each missing pathway, and then calculates a test statistic, Fisher’s *C*, using the following equation: *C* = –2 × ln (*P_i_*), where *P_i_* is the *P*-value of the *i*-th missing pathway. Fisher’s *C* can then be compared to a chi-square distribution with 2*k* degrees of freedom, where *k* is the total number of missing pathways. If there are many missing pathways with low *P*-values, this will result in a lower *P*-value for the structural equation model. Therefore, a *P*-value < 0.05 indicates a poor fit for the structural equation model, whereas a *P*-value > 0.05 indicates a good fit. Note that if we have included the key plant traits as well as biotic and abiotic factors, then there should be no missing paths between community responses and willow genotype or our environmental treatments. If we identified a missing path due to willow genotype, we included genotype averages from principle components of willow phenolic chemistry (salicylate/tannin PC1, phenolic acid PC1-2, flavonoid PC1-2, miscellaneous flavonoids PC1) measured in our prior study (Barbour *et al.* 2015) as additional predictors in the piecewise GLMMs. Details on the principal components analysis of phenolic compounds are given in Barbour *et al.* (2015).

All analyses were conducted in R version 3.2.4 (R Core Team 2016). All data and R-scripts for reproducing the reported results are publicly available and can be found at the GitHub repository https://github.com/mabarbour/Lanphere_Experiments.git. The release associated with this version of the manuscript can be found at https://doi.org/10.5281/zenodo.840033.

## Results

### How do willow genotype and the environment affect plant functional traits and other environmental factors?

#### Ant-aphid experiment

We hypothesized that the effect of willow genetic variation and the biotic environment on arthropod communities would be mediated, in part, by variation in the abundance of *A. farinosa* and *F. obscuripes.* While distance from ant mounds had little effect on *A. farinosa* (*χ*_1_ = 0.55, *P* = 0.460), willow genotype had a strong effect, with the average number of aphids ranging from 0.05 to 7 individuals among the most disparate willow genotypes in the aphid treatment (*χ*_9_ = 20.83, *P* = 0.013). This strong effect of willow genotype on *A. farinosa* in the aphid treatment resulted in a G×E_aphid_ effect on the abundance of *F. obscuripes* (*χ*_2_ = 6.26, *P* = 0.044), with ant abundance varying from 0 to ∼0.5 individuals (on average) among clones in the aphid treatment, whereas they were virtually absent when aphids were excluded. Proximity to ant mounds had no effect on the abundance of *F. obscuripes* (*χ*_1_ = 1.68, *P* = 0.195).

In addition to ant-aphid interactions, we hypothesized that the effect of willow genetic variation and the biotic environment on arthropod communities would be mediated by plant traits. We observed both direct and interactive effects of willow genotype and the biotic environment on plant traits (Table A1 in Supporting Information). All of the plant-growth traits we measured varied approximately 2-fold among the most disparate willow genotypes (plant height *F*_9,204.2_ = 15.83, *P* < 0.001; shoot count *χ*_9_ = 65.84, *P* < 0.001; shoot length *F*_9,204.2_ = 7.27, *P* < 0.001). Willows did appear to produce 28% more shoots in the absence of aphids, but only at the furthest distance from ant mounds (E_aphid_ × E_ant_ *χ*_1_ = 4.20, *P =* 0.040). While there was little apparent effect of willow genotype and the biotic environment on leaf water content (Table A1), we found that the addition of aphids modified the effect of certain willow genotypes on leaf trichome density (G×E_aphid_ *χ*_8_ = 23.17, *P* = 0.003). Specifically, two clones (S and T) produced ∼4-fold more trichomes when aphids were absent, whereas genotype L produced 3-fold more trichomes when aphids were present (Table A1).

#### Wind experiment

One of the mechanisms by which wind exposure could influence willow-associated communities is through accumulated effects on soil properties; however, we observed only modest effects of wind exposure on soil properties (Table A2). Specifically, soil in wind-exposed plots was marginally drier (*F*_1,9.0_ = 3.52, *P* = 0.093) with higher amounts of total Nitrogen (*F*_1,9.0_ = 3.52, = 5.08, *P* = 0.051) than in unexposed plots, but there was no clear difference in either percent organic matter (*F*_1,8.4_ = 0.68, *P* = 0.434) or nutrient PC1 (*F*_1,9.0_ = 1.31, *P* = 0.282).

As with the ant-aphid experiment, we hypothesized that the effects of wind exposure and willow genotype on associated communities would be mediated by plant traits. Interestingly, we found that plant-growth and leaf quality traits responded differently to wind exposure and willow genetic variation (Table A2). For example, wind exposure negatively affected all plant-growth traits (plant height *F*_1,9.0_ = 29.10, *P* < 0.001; shoot count *χ*_1_ = 9.91, *P* = 0.002; shoot length *F*_1,9.0_ = 10.44, *P* = 0.010). Moreover, the negative effects of wind exposure were magnified by the end of the experiment for both plant height (*F*_1,158.4_ = 16.69, *P* < 0.001) and the number of shoots produced (*χ*_1_ = 12.53, *P* < 0.001). Still, willow genotype had a pronounced effect on all plant-growth traits, resulting in willows that varied over 2-fold in height (*F*_9,145.3_ = 9.13, *P* < 0.001), number of shoots (*χ*_9_ = 47.42, *P* < 0.001), and shoot length (*F*_9,144.2_ = 4.97, *P* < 0.001) among the most disparate genotypes. While the effect of willow genotype on the number of shoots changed by the end of the experiment (*χ*_9_ = 18.26, *P* = 0.032), this G×E_year_ effect was relatively small (*R*^2^ = 0.05) compared to the effect of genotype alone (*R*^2^ = 0.13). In contrast to plant growth, willow genotype was the primary factor in determining leaf quality traits across both years of the experiment (Table A2). The leaves of willow genotypes varied 46-fold in trichome density (*χ*_9_ = 67.31, *P* < 0.001), 1.5-fold in SLA (*F*_9,122.5_ = 4.21, *P* < 0.001), and 1.6-fold in C:N (*F*_9,70.5_ = 4.88, *P* < 0.001). We had data available on leaf water content for 2012 and 2013, and we found that the amount of variation explained by willow genotype depended on the sampling year (*F*_9,141.6_ = 2.80, *P* = 0.005; 2012, *R*^2^ = 0.11; 2013, *R*^2^ = 0.16). Unlike aboveground plant traits, root C:N did not appear to be influenced by either wind exposure (*F*_1,8.7_ = 0.31, *P* = 0.590) or willow genotype (*F*_9,107.0_ = 0.85, *P* = 0.569).

### How do willow genotype and the environment shape the diversity and composition of above and belowground communities?

#### Ant-aphid experiment

Willow genotype tended to be more important than the biotic environment in structuring the arthropod community (Table A1). We found that average arthropod richness varied from 1.2 to 3.2 species among genotypes (*σ*^2^ = 11%, *χ*_9_ = 41.35, *P* < 0.001), while arthropod abundance varied 4-fold among the different clones (Fig. 1A; *σ*^2^ = 7%, *χ*_9_ = 34.86, *P* < 0.001). The effect of willow genotype on arthropod richness was explained by correlated responses in arthropod abundance, as there was no difference in rarefied richness among genotypes (*F*_9,138.8_ = 0.83, *P* = 0.586). Aphid treatment was the only factor that affected rarefied richness (*σ*^2^ = 4%, *F*_1,139.2_ = 5.34, *P* = 0.022), resulting in a 16% decrease in rarefied richness when aphids were added to willows (Fig. 1C); however, this effect of aphid treatment did not translate into an effect on total richness (*χ*_1_ = 0.45, *P* = 0.504). Willows in the aphid treatment also had 2-fold more arthropods, but only at the furthest distance from ant mounds (Fig. 1B, *σ*^2^ = 9%, *χ*_1_ = 8.12, *P* = 0.004). Proximity to ant mounds did not influence any other aspect of the arthropod community (Table A1). In contrast to richness and abundance responses, arthropod community composition was influenced by an interaction between willow genotype and the aphid treatment (Fig. 2A; 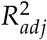 = 6%, *F*_9,157_ = 1.45, *P* = 0.022). This G×E effect was primarily due to the differential response of other aphids (Table A3) to a single willow genotype (Fig. 2A solid line). If we remove this genotype from the analysis, we find independent effects on community composition, with the effect of willow genotype (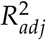 = 3%, *F*_8,156_ = 1.66, *P* = 0.007) being relatively more important than the addition of aphids (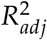 = 1%, *F*_1,156_ = 2.93, *P* = 0.017).

**Figure 1:**
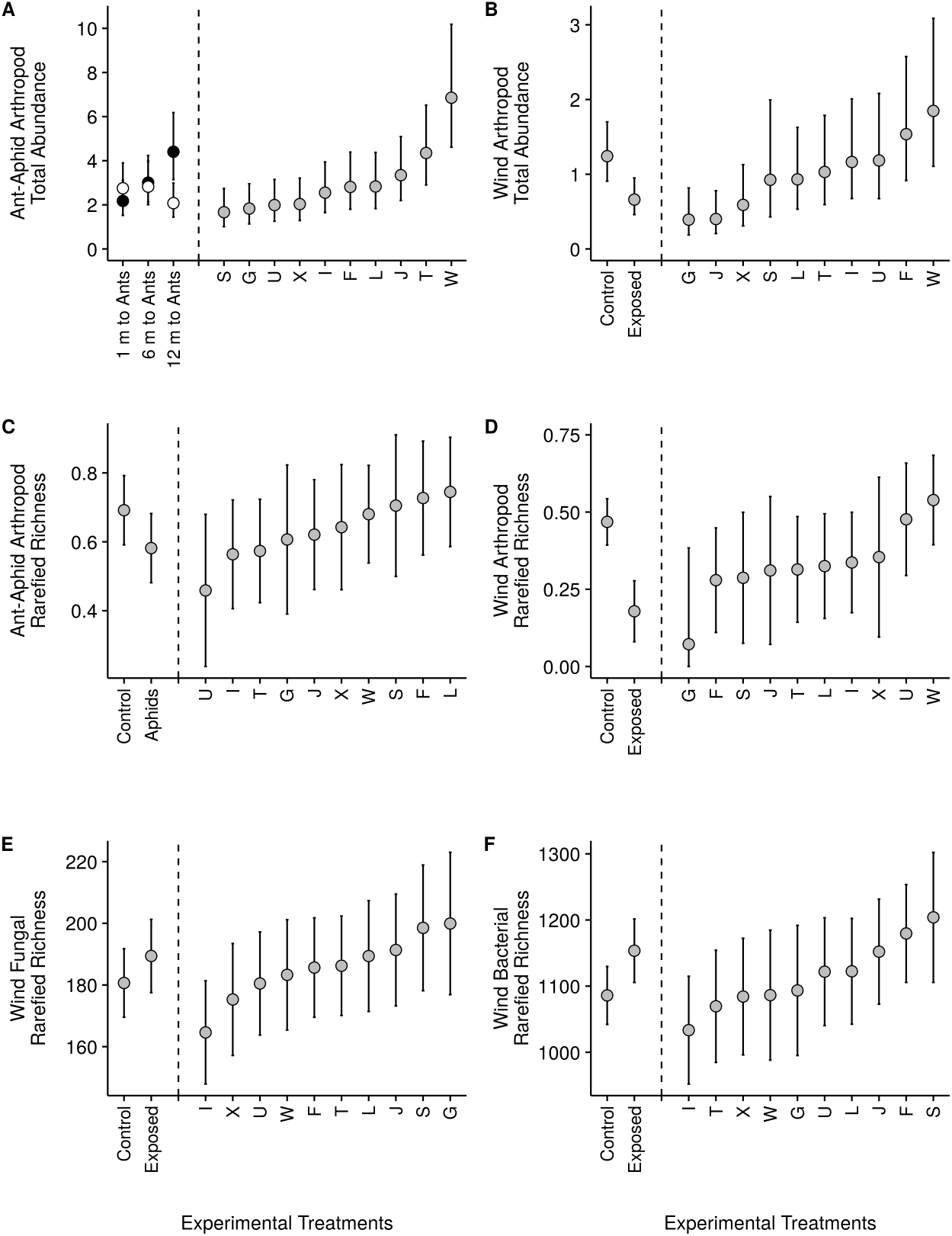
Effects of genetic variation within the willow *Salix hookeriana* as well as biotic (aphid additions and proximity to ant mounds) and abiotic (wind exposure) factors on the structure of above and belowground communities. Willow genotype had strong effects on arthropod abundance in both the ant-aphid (A) and wind exposure (B) experiments. In the ant-aphid experiment, arthropod abundance was influenced by the addition of the aphid *Aphis farinosa*, but only at the furthest distance from mounds of the ant *Formica obscuripes* (A). In the wind experiment, wind exposure reduced arthropod abundance (B). In contrast to arthropod abundance, willow genotype had weak effects on arthropod rarefied richness in both the ant-aphid (C) and wind exposure (D) experiments. In the ant-aphid experiment, the addition of aphids reduced the probability of encountering a different arthropod species (rarefied richness, C). In the wind experiment, wind exposure dramatically reduced arthropod rarefied richness (D). Similar to arthropod rarefied richness, willow genotype had weak effects on the rarefied richness of rhizosphere fungi (E) and bacteria (F). While wind exposure had a weak effect on fungal rarefied richness (E), we found that wind exposed willows hosted a more diverse community of rhizosphere bacteria (F). Points and error bars correspond to the response variable’s mean ± 95% confidence interval. We calculated mean and confidence intervals based on the full models (Tables A1,A4) using the *effects* package in R.

**Figure 2:**
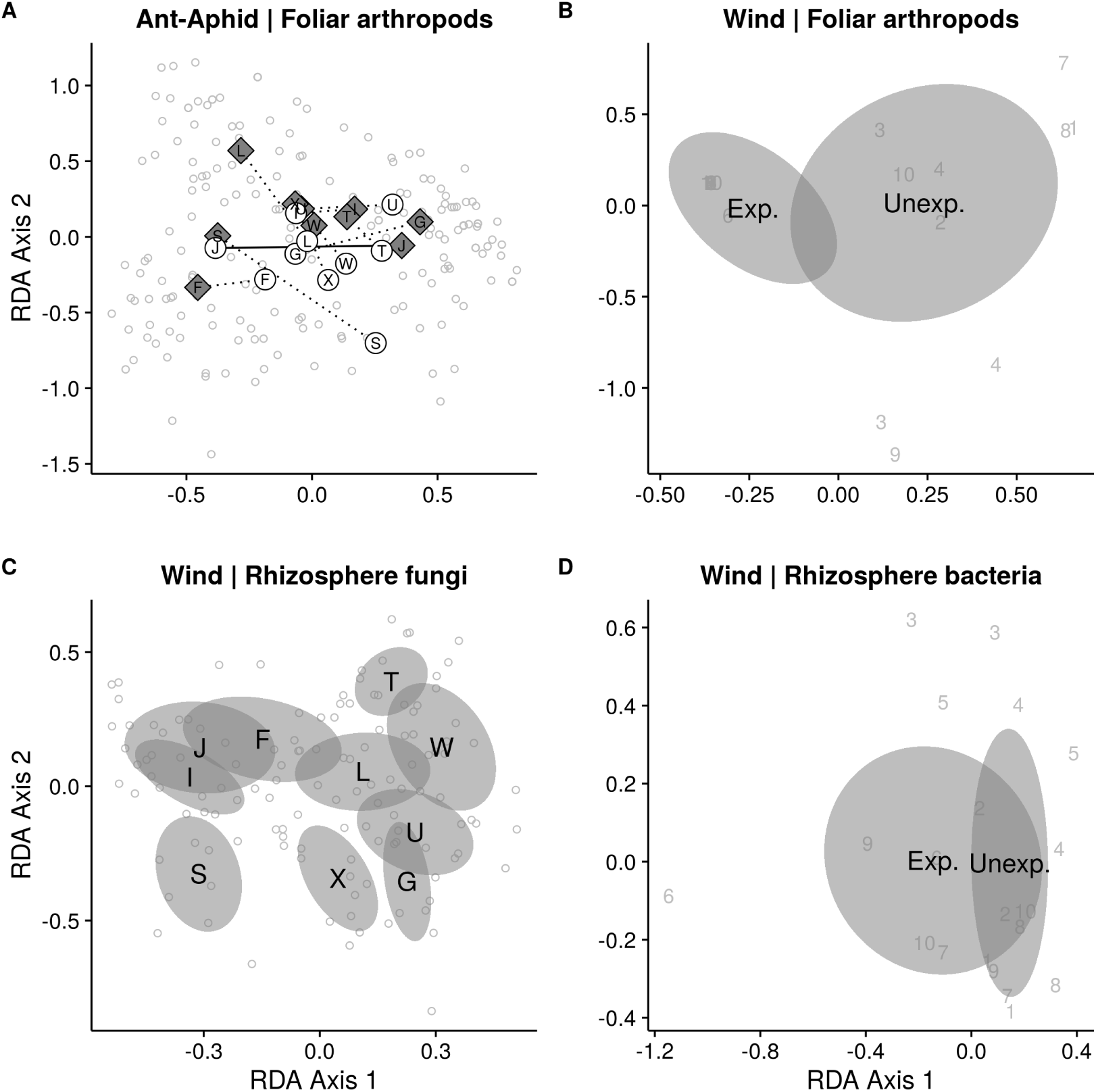
Effects of genetic variation within the willow *Salix hookeriana* as well as biotic (aphid additions) and abiotic (wind exposure) factors on the composition of above and belowground communities. In the ant-aphid experiment (A), the addition of the aphid *Aphis farinosa* (grey diamonds) modified the effect of willow genotype on the composition of the arthropod community. This interaction between willow genotype and aphid treatment was solely due to the differential effect of genotype J (solid line) on the abundance of non-*A. farinosa* aphids in the aphid treatment (table A3). For arthropods in the wind experiment (B), we found that wind exposure had a strong effect on community composition in 2013 (but not 2012, table A4), with no clear effect of willow genotype. Belowground, we found that willow genotype influenced the composition of rhizosphere fungi (C), while the composition of rhizosphere bacteria was only influenced by wind exposure (D). Black text and grey ellipses correspond to the community centroid ± 95% confidence interval. Grey numbers denote blocks and each unique number is the community centroid for the plot within each block. Grey circles mark the location of individual willow communities in multivariate space. We calculated the locations of centroids ± 95% confidence interval and individual samples using redundancy analysis on Hellinger-transformed community data.

#### Wind experiment

In contrast to the ant-aphid experiment, we found that the abiotic environment tended to be more important than willow genotype in structuring the arthropod community (Table A4). In particular, willows growing in wind-exposed plots hosted 51% fewer species (*σ*^2^ = 11%, *χ*_1_ = 13.55, *P* < 0.001), 47% fewer individuals (Fig. 1B; *σ*^2^ = 2%, *χ*_1_ = 5.48, *P* = 0.019), and 60% fewer rarefied species (Fig. 1D; *σ*^2^ = 23%, *F*_1,7.8_ = 22.82, *P* = 0.001) compared to unexposed willows. In spite of the effects of wind exposure, willow genotype had clear effects on both the richness (∼3-fold differences, *σ*^2^ = 6%, *χ*_9_ = 28.01, *P* < 0.001) and abundance (Fig. 1B; ∼5-fold differences, *σ*^2^ = 5%, *χ*_9_ = 25.25, *P* = 0.003) of arthropods, but only a marginal effect on rarefied richness (Fig. 1D; *σ*^2^ = 5%, *F*_9,71.1_ = 1.96, *P* = 0.058). Arthropod communities on willows had both more species (*σ*^2^ = 4%, *χ*_1_ = 10.33, *P* = 0.001) and more individuals (*σ*_2_ = 2%, *χ*_1_ = 6.72, *P* = 0.010) in the second year of the experiment compared to the first; however, we also conducted more arthropod surveys for the wind experiment in 2013 vs. 2012. In terms of community composition, we observed strong effects of wind exposure by the end of experiment (Fig. 2B; 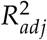 = 40%, *F*_1,68_ = 12.80, *P* = 0.001). These compositional differences were driven primarily by gall midges (Family: Cecidomyiidae) being relatively less abundant on wind-exposed willows (*χ*_1_ = 16.28, *P* < 0.001), whereas leaf-tiering moths (Family: Tortricidae) were insensitive to wind exposure (and therefore relatively more abundant; *χ*_1_ = 1.34, *P* > 0.05). Although several arthropod taxa varied in total abundance among willow genotypes (Table A5), we did not detect an effect of genotype on community composition in either year of the experiment (2012: *F*_9,51_ = 0.96, *P* = 0.502; 2013: *F*_9,68_ = 1.17, *P* = 0.271).

Willow genotype and wind exposure had distinct effects on root-associated fungal and bacterial communities compared to foliar arthropods (Table A4). Neither wind exposure (*F*_1,8.8_ = 0.90, P = 0.369) nor willow genotype (*F*_9,95.0_ = 1.21, *P* = 0.295) influenced the rarefied richness of fungal OTUs (Fig. 1E). However, the composition of the fungal community did vary among willow genotypes (Fig. 1C; 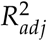 = 1%, *F*_9,117_ = 1.00, *P* = 0.005) with no detectable effect of wind-exposure (*F*_1,9_ = 1.19, *P* = 0.162). Genotypic differences in fungal community composition were due to variation in the relative abundance of at least eight different OTUs, most of which were saprotrophs (Table A6) and not arbuscular mycorrhizae or ectomycorrhizae.

In contrast to the fungal community, wind exposure influenced the bacterial community (Table A4), but in the opposite direction of foliar arthropods. For example, the rarefied richness of bacterial OTUs was 6% higher on the roots of wind-exposed vs. unexposed plants (Fig. 1F; *σ*^2^ = 7%, *F*_1,7.9_ = 6.52, *P* = 0.034). While wind exposure had a marginal effect on the composition of the bacteria community (Fig. 1D; 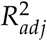 = 3%, *F*_1,9_ = 1.38, *P* = 0.092), there was no detectable effect of willow genotype on any aspect of the bacterial community (rarefied richness: *F*_9,100.3_ = 1.63, *P* = 0.117; community composition: *F*_9,120_ = 0.93, *P* = 0.536).

#### What is the relative importance of genetic variation, phenotypic plasticity, and direct environmental effects in structuring communities?

##### Ant-aphid experiment

Using structural equation models (for richness, abundance, and rarefied richness) and redundancy analysis (for community composition), we found that genetic variation in plant traits and direct effects of aphid additions were the primary determinants of the arthropod community rather than phenotypic plasticity (Table 1). The effect of genetic variation on arthropod richness and abundance was mediated primarily by plant trait PC1 (Fig. 3A,B). Plant height, shoot count, and shoot length all had strong, positive loadings on trait PC1 (Table A7), indicating that larger willows hosted more arthropod species and individuals. Arthropod abundance was also positively influenced by the addition of aphids, primarily because *A. farinosa* attracted individuals of other ant species (Pearson’s *r* = 0.42, *t*_282_ = 7.74, *P* < 0.001) and these other ants were the second most abundant taxonomic group in the community. In contrast to total abundance, the addition of aphids negatively affected rarefied richness (Fig. 3C). This negative effect was due in part to aphid additions attracting more *F. obscuripes*, an active generalist predator that likely consumed or inhibited the colonization of other arthropods. In terms of composition, we found that the abundance of *A. farinosa* was the only factor (of the mechanisms we modeled) influencing the arthropod community (*F*_1,183_ = 2.86, *P* = 0.025). Specifically, higher abundance of *A. farinosa* resulted in an increase in the proportional abundance of other ant species in the community (Fig. 3D).

**Table 1:**
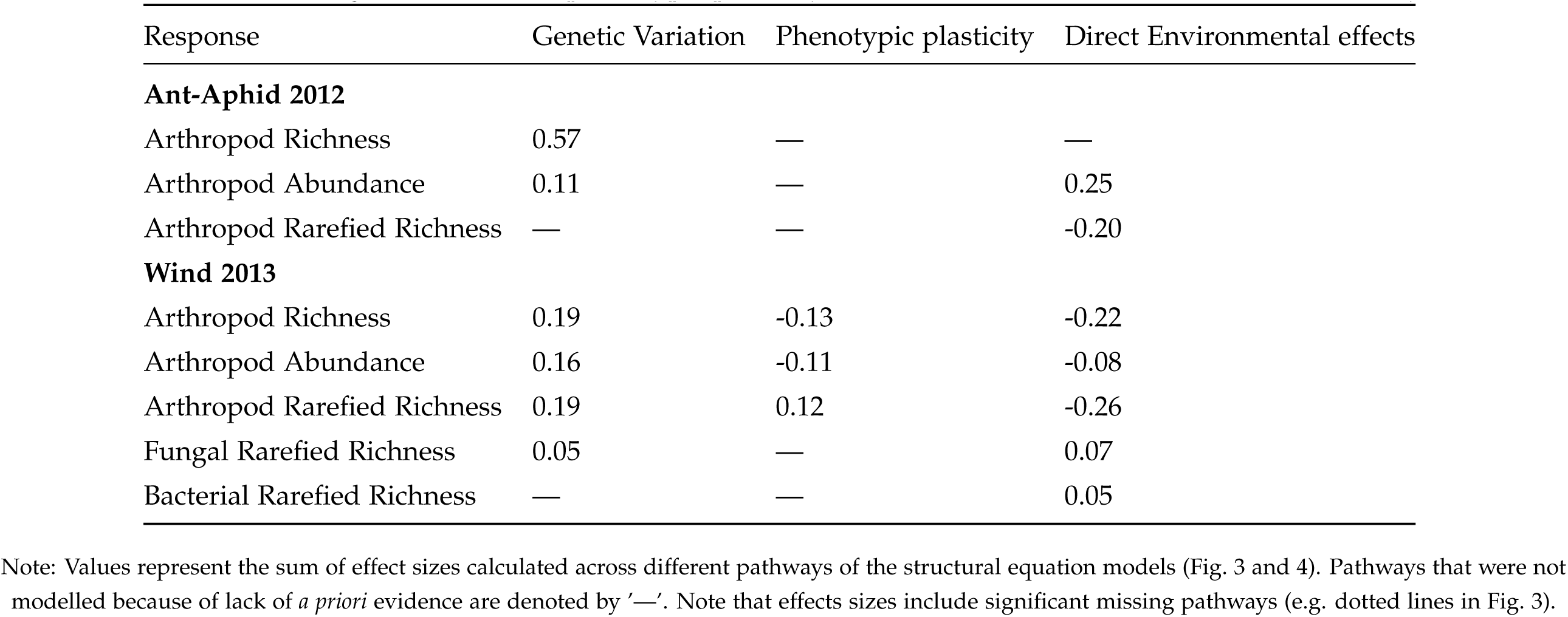
Net effect sizes of genetic variation, phenotypic plasticity, and direct environmental effects on community structure.

**Figure 3:**
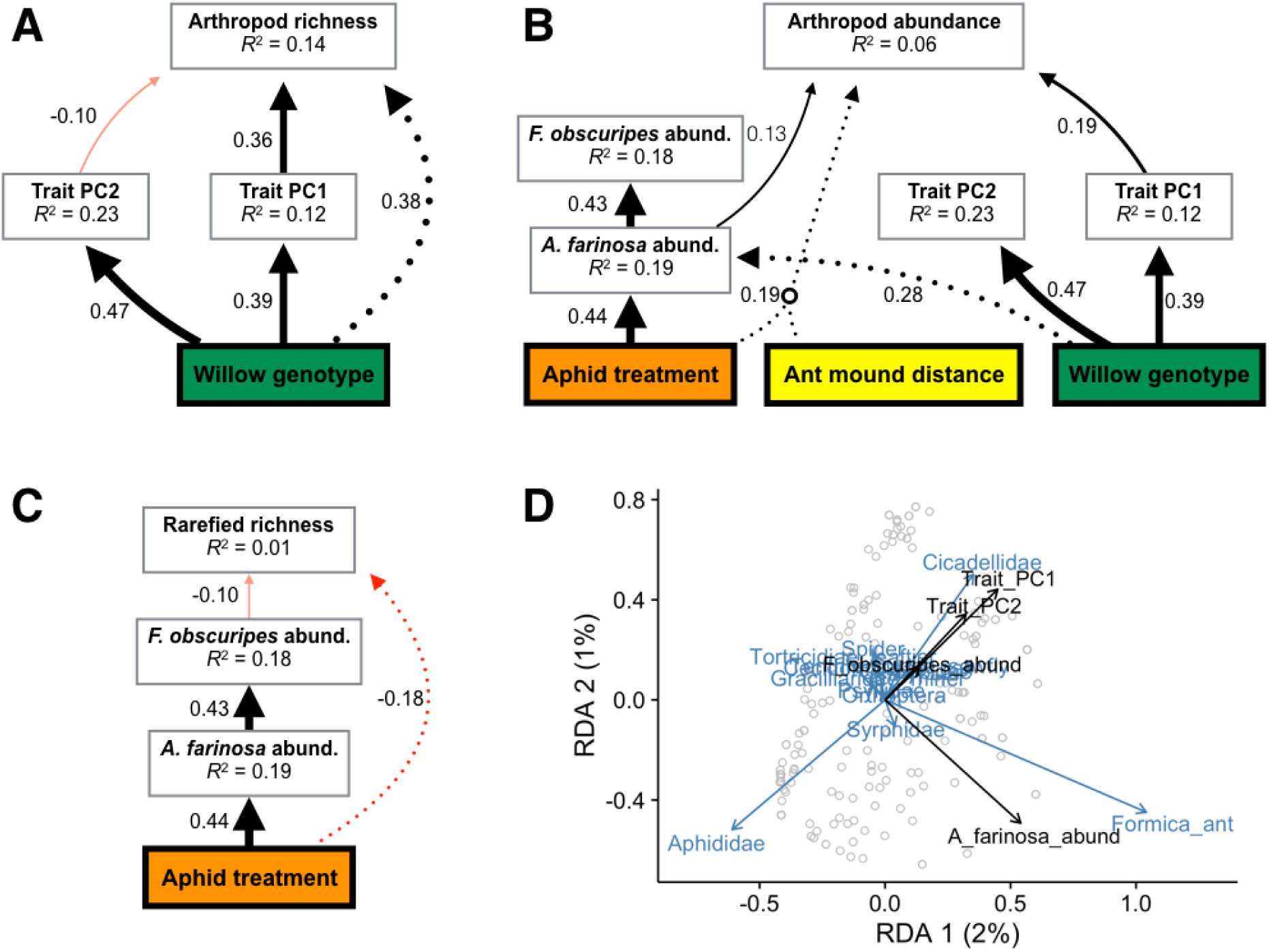
Statistical models of the processes mediating arthropod community assembly in the ant-aphid experiment. Piecewise structural equation models of arthropod richness (A), abundance (B), and rarefied richness (C). Colored and white boxes represent exogenous and endogenous variables, respectively. Solid arrows correspond to modeled pathways between predictor and response variables, and may be either positive (black) or negative (red), while dotted arrows correspond to missing paths. For clarity, we only plotted paths with standardized coefficients > 0.10, but marginally significant effects (0.05 < P < 0.10) are transparent. Numbers next to all arrows represent the standardized path coefficient, which also corresponds to the thickness of arrows. (D) Redundancy analysis illustrating the effect of plant traits (Trait PC1, PC2) and *A. farinosa* abundance on the arthropod community (Hellinger-transformed = square root of proportional abundances of species found on each willow). Black and blue arrows correspond to plant traits and species, respectively, while grey dots represent the position of individual willow communities.

Despite our detailed analysis of potential mechanisms, our structural equation models revealed multiple missing paths (dotted lines, Fig. 3A,B,C), resulting in rather poor fits for most models (richness: *C*_2_ = 24.84, *P* < 0.001; abundance: *C*_32_ = 48.88, *P* = 0.029; rarefied richness: *C*_4_ = 10.52, *P* = 0.032). For example, after accounting for the traits we measured, willow genotype still had a strong effect on arthropod richness (Fig. 3A) and *A. farinosa* abundance (Fig. 3B). These genotypic effects are likely due to heritable variation in willow leaf chemistry, given the strong effects we observed of phenolic acid PC1 (richness *F*_1,266.13_ = 22.71, *P* < 0.001; *A. farinosa F*_1,263.72_ = 6.73, *P* = 0.010), flavonoids PC2 (*A. farinosa F*_1,264.81_ = 4.14, *P* = 0.043), and miscellaneous flavonoids PC1 (richness *F*_1,269.94_ = 15.51, *P* < 0.001) when we included these traits as predictors in our analysis. While we expected the effects of our aphid treatment to be mediated by aphid abundance, the missing paths between aphid treatment and arthropod abundance (Fig. 3B) as well as rarefied richness (Fig. 3C) suggests that the presence of aphids alone influenced arthropod community responses. From our redundancy analysis, we found that *A. farinosa* abundance was an important determinant of community composition (*F*_1,183_ = 2.86, *P* = 0.021), but we still failed to detect the G×E_aphid_ effect (*F*_9,164_ = 1.71, *P* = 0.004). However, if we account for known heritable variation in leaf flavonoid chemistry, then we can explain the G × E_aphid_ effect on the arthropod community (flavonoid PC1×E_aphid_ *F*_1,178_ = 5.20, *P* = 0.001; miscellaneous flavonoids PC1×E_aphid_ *F*_1,178_ = 2.58, *P* = 0.020).

##### Wind experiment

Similar to the ant-aphid experiment, we found that genetic variation and direct effects of wind exposure were the primary factors influencing the arthropod community rather than phenotypic plasticity (Table 1). Both trait PC1 and PC2 mediated the indirect effects of willow genetic variation and wind exposure (negative) on the arthropod community (Fig. 4A), but the effects of genetic variation were relatively stronger (Table 11). Trait PC1 had a strong, positive effect on arthropod richness (*β* = 0.37), abundance (*β* = 0.28), and rarefied richness (*β* = 0.37, Fig. 4A). Similar to the ant-aphid experiment, trait PC1 had strong, positive associations with plant height, shoot count, and shoot length (Table A7), indicating that larger willows hosted more arthropod individuals and species. Trait PC2 had a smaller, but negative effect on arthropod richness (*β* = –0.13), abundance (*β* = –0.15), and rarefied richness (*β* = –0.12, Fig. 4A). Trait PC2 had a strong positive correlation with leaf C:N, but strong negative correlations with leaf water content and SLA (Table A7), indicating that willows with poorer quality leaf tissue hosted fewer arthropod individuals and species. Although the community-level effects of wind exposure due to phenotypic plasticity were relatively weak, wind exposure had strong and direct, negative effects on arthropod richness (*β* = –0.22), abundance (*β* = –0.08), and rarefied richness (*β* = –0.26, Fig. 4A). The qualitative effects of genetic variation, phenotypic plasticity, and direct environmental effects held for the richness, abundance, and rarefied richness of foliar arthropods in the first year of the experiment as well (*C*_22_ = 28.70, *P* = 0.154), except that trait PC2 was determined by different traits (Table A7) and did not appear to affect any aspect of the arthropod community (richness, *P* = 0.657; abundance, *P* = 0.104; rarefied richness, *P* = 0.850). For community composition, we only analyzed the data from the second year of the experiment because this was the only year for which we detected a significant effect of wind exposure. We found that the effects of wind exposure on community composition were primarily mediated by plant trait PC1 (*F*_1,76_ = 12.05, *P* = 0.001). Positive values of trait PC1 (i.e. larger plants) had greater proportional abundance of gall midges, leaf-mining moths, and spiders, whereas leaf-tiering moths were insensitive to plant size (Fig. 4B).

**Figure 4:**
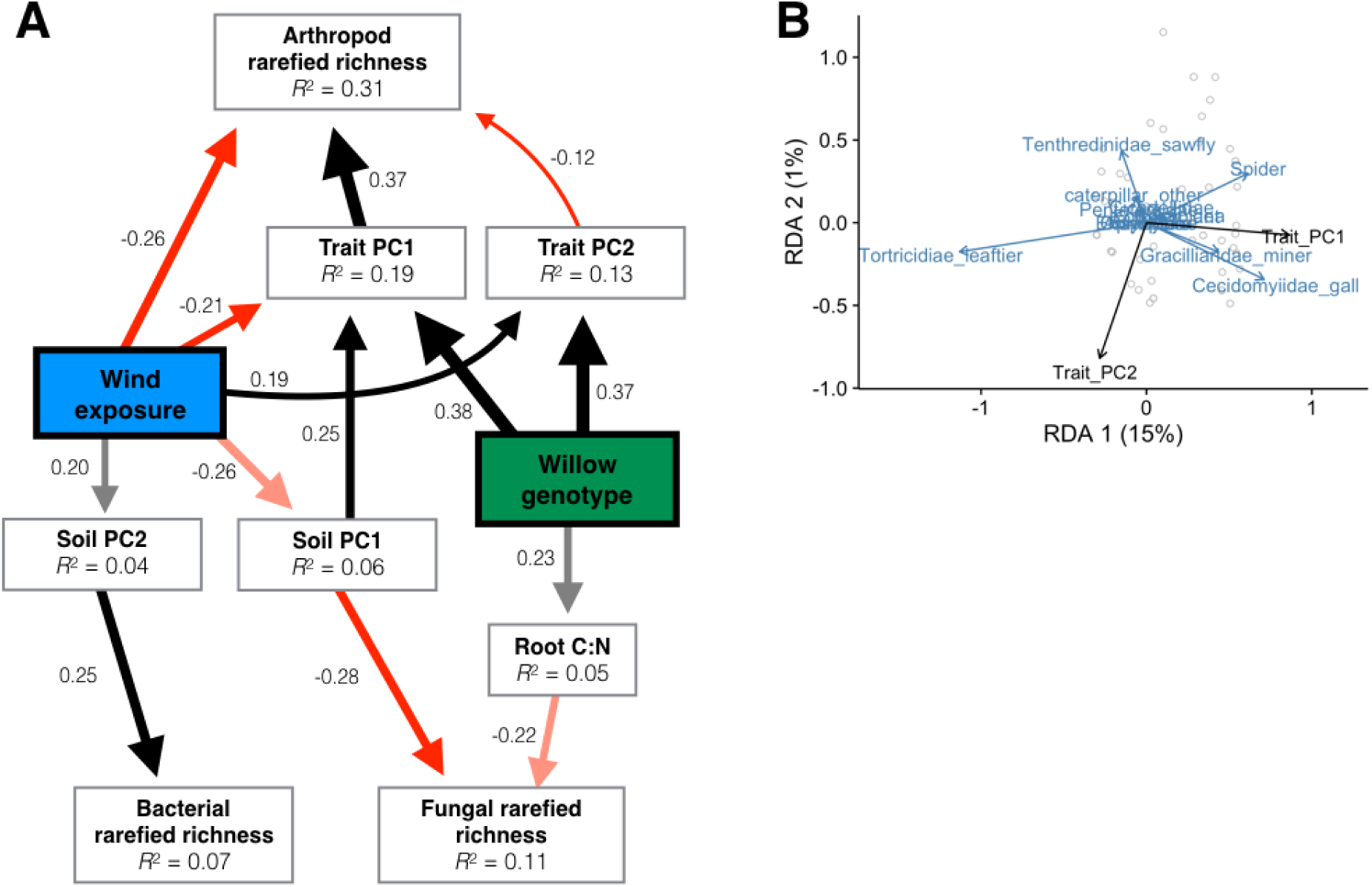
Statistical models of the processes mediating community assembly in the wind experiment for 2013. (A) Piecewise structural equation model of the rarefied richness of foliar arthropods as well as rhizosphere fungi and bacteria. Colored and white boxes represent exogenous and endogenous variables, respectively. Solid arrows correspond to modeled pathways between predictor and response variables, and may be either positive (black) or negative (red). For clarity, we only plotted paths with standardized coefficients > 0.10, but marginally significant effects (0.05 < P < 0.10) are transparent. Numbers next to all arrows represent the standardized path coefficient, which also corresponds to the thickness of arrows. (B) Redundancy analysis illustrating the effect of plant traits (Trait PC1, PC2) on the arthropod community (Hellinger-transformed = square root of proportional abundances of species found on each willow). Black and blue arrows correspond to plant traits and species, respectively, while grey dots represent the position of individual willow communities.

As with the aboveground community, we found that genetic variation and direct environmental effects were the primary determinants of rhizosphere communities, although we did a poorer job of identifying the specific mechanisms. For example, we previously showed that fungal community composition varied among willow genotypes (Table A4), but we failed to identify the root traits mediating the effect of willow genetic variation (*F*_9,106_ = 1.03, *P* = 0.002). Interestingly though, we found that heritable variation in leaf phenolic acid PC1 (*F*_1,112_ = 1.11, *P* = 0.062), flavonoid PC1 (*F*_1,112_ = 1.36, *P* = 0.001), and miscellaneous flavonoids PC1 (*F*_1,112_ = 1.15, *P* = 0.028) could account for willow genetic effects on fungal community composition. While wind exposure had only a modest effect on soil properties, soil characteristics had strong direct effects on the rarefied richness of rhizosphere communities. For example, soil PC1 (*β* = –0.28), and to a lesser extent root C:N (*β* = –0.22), negatively affected fungal rarefied richness (Fig. 4A). Soil PC1 had strong positive correlations with soil moisture and organic matter, but negative correlations with 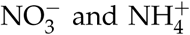 (Table A8), indicating that fungal communities were more diverse in drier environments with more available nitrogen. In contrast, soil PC2 was the primary factor in determining bacterial rarefied richness (*β* = 0.25). Micronutrients such as Ca^2+^, Mg^2+^, and Cd^2+^ had strong positive loadings on soil PC2 (A8), indicating that bacterial richness was greater in environments with more of these micronutrients. Although we detected clear effects of soil properties and root C:N on rarefied richness of rhizosphere communities, none of these soil and plant characteristics were strong predictors of community composition (Table A9).

In contrast to the ant-aphid experiment, our structural equation models provided good fits to our data (i.e., *P* > 0.05), indicating that we identified the key processes affecting the richness (*C*_10_ = 2.94, *P* = 0.983), abundance (*C*_10_ = 6.73, *P* = 0.751), and rarefied richness (Fig. 4A; *C*_32_ = 23.32, *P* = 0.899) of willow-associated communities.

## Discussion

### How do willow genotype and the environment affect plant functional traits and other environmental factors?

Across both environmental gradients, willow genotype had clear effects on plant growth, but usually a stronger influence on leaf quality traits. This result corresponds with the generally high heritability of leaf quality compared to plant growth traits we observed in our prior work with *Salix hookeriana* (Barbour *et al.* 2015) and others have observed in other plant species (Geber & Griffen 2003; Johnson *et al.* 2009). These differences in heritable trait variation predict that plant growth traits will be more influenced by environmental variation than leaf quality. In correspondence with this prediction, we found that ant-aphid interactions and wind exposure minimally influenced the expression of leaf quality traits. As predicted, wind exposure had a strong negative effect on plant growth, but ant-aphid interactions had only weak and idiosyncratic effects. These results suggest that the plasticity of plant traits depends not only on plant genotype, but also on the type of plant trait and environmental factor. Since the assembly of host-plant communities is often influenced by multiple plant traits that vary in their degree of heritability (Agrawal 2005; Barbour *et al.* 2015), our work suggests that predicting the community assembly on host plants in different environments will be a challenging task.

The phenotypic effects of willow genotype were usually more important than the environment in both experiments. This result, however, could be biased due to the relatively short time scale of this experiment. Indeed, the effects of wind exposure on plant growth were increasing by the end of the experiment, indicating that wind exposure may have become more important than willow genotype if we ran the experiment longer. The relatively short time scale of this experiment may also explain why we only observed weak and idiosyncratic G × E effects on plant traits. In addition, our use of stem cuttings may have given the effect of willow genotype an initial advantage in the different environments. While transplanting greenhouse-grown plants to the field is common in community genetics studies, this minimizes the effects of environmental variation on phenotypic plasticity. We suggest that future genotype-by-environment studies account for the effects of environmental variation throughout plant development to more accurately characterize the effects of phenotypic plasticity.

### How do willow genotype and the environment affect the diversity and composition of above and belowground communities?

The relative importance of host-plant genetic effects on its associated arthropod community depended on the type of environmental gradient. In the ant-aphid experiment, willow genetic variation tended to have a stronger effect on arthropod community structure compared to aphid additions and proximity to ant mounds. This result supports an emerging generalization that host-plant genetic variation is often more important than the presence of arthropod herbivores (Cronin & Abrahamson 2001; Fritz *et al.* 1986; Johnson 2008; McGuire & Johnson 2006) or predatory ants (Abdala-Roberts *et al.* 2012; Johnson 2008; Mooney & Agrawal 2008) in structuring arthropod communities. This is likely due to host plants providing all of the habitat for the associated communities, compared to the smaller spatial scale at which ant-aphid interactions occur. Still, we did find that aphid additions reduced arthropod diversity (rarefied richness) and modified the effect of willow genotype on the composition of the arthropod community, suggesting that accurately predicting community assembly requires an understanding of both factors.

In the wind experiment, we found that wind exposure dominated willow genotype in the strength of its effect on foliar arthropods, supporting the notion that wind is a key environmental factor in structuring communities associated with host plants in coastal dune ecosystems (Crutsinger *et al.* 2014, 2010; Miller & Weis 1999). Similarly, Crutsinger *et al.* (2014) found that there were more arthropod species and individuals on prostrate vs. erect morphs of coyote bush (*Baccharis piluaris*), due to their low-lying growth form which enabled them to be more productive than erect morphs at their windy coastal dunes site. Unfortunately, few studies have examined community-level responses to natural variability in specific abiotic factors and host-plant genetic variation, making it difficult to draw other useful comparisons. The majority of genotype-by-abiotic environment studies to date have used fertilizers to manipulate soil nutrient availability (Abdala-Roberts *et al.* 2012; Barrios-Garcia *et al.* 2016; Orians & Fritz 1996; Rossi & Stiling 1998), but it is unclear whether these manipulations reflect natural variation in soil nutrients. This may explain why the effects of soil nutrient variability range from being independent and weak (Abdala-Roberts *et al.* 2012; Barrios-Garcia *et al.* 2016) to being strong modifiers (Orians & Fritz 1996) of host-plant genotype on arthropod communities. The only other genotype-by-abiotic environment study that we are aware of manipulated sun exposure to sea daisies (*Borrichia frutescens*, Rossi & Stiling 1998) and, similar to our study, observed strong, independent effects of sun exposure on densities of the gall midge, *Asphondylia borrichiae.* If we are to make progress on understanding the relative importance of willow genotype vs. the environment for associated communities, future experimental work should focus on manipulating natural variation in specific abiotic factors, or across natural gradients in the abiotic environment.

Although diverse assemblages of above and belowground taxa colonize host plants, only one other genotype-by-environment study, to our knowledge, has simultaneously measured the responses of above and belowground assemblages in different environments (Wagner *et al.* 2016). We found that foliar arthropods responded differently than rhizosphere fungi and bacteria to wind exposure and willow genetic variation, suggesting that these communities are responding to different plant traits and environmental correlates of wind exposure. This lack of correspondence between above and belowground community responses has been observed in other studies (Lamit *et al.* 2015; Wagner *et al.* 2016). Although Crutsinger *et al.* (2014) found similar responses in above and belowground communities, they also found that different mechanisms mediated the assembly of these communities. Future work with belowground communities should strive to include more comprehensive measurements of root traits to get a better understanding of the mechanisms mediating the assembly of these communities. Understanding associations between above and belowground traits will be important for predicting when we would expect linkages between above and belowground communities.

### What is the relative importance of genetic variation, phenotypic plasticity, and direct environmental effects in structuring communities?

Across both experiments, we found that the effects of host plant traits on foliar arthropod communities were primarily mediated by genetic variation rather than phenotypic plasticity. Although the indirect effect of environmental variation was of minor importance, we found that the biotic and abiotic environment could have strong direct effects on community structure.

While willow genotype had a strong effect on leaf quality traits, the effects of genetic variation on arthropod community structure were primarily mediated by plant size in both experiments. We found similar results in our prior work with *Salix hookeriana* (Barbour *et al.* 2015), although the effects of plant size appeared much more important in this study. This is likely because the absolute size of the plants in the current study were small (<60 cm tall), therefore size could be a limiting resource in determining arthropod community structure. Genotype-by-environment experiments with small-statured plants (forbs, herbs, etc.) have similarly found that plant size is a key factor in structuring arthropod communities (Crutsinger *et al.* 2014; Johnson & Agrawal 2005). Therefore, the effects of host-plant size on arthropod communities may be especially strong for smaller plants, with leaf quality traits becoming more important as plants grow. While the importance of plant size corresponds with other work in coastal dunes (Crutsinger *et al.* 2014), studies in ant-aphid systems often find that the community-level effects of genetic variation are mediated by ant abundance. One possible reason for this discrepancy is that the absolute variation in *F. obscuripes* abundance was quite low in our experiment (max. genotype average = ∼0.5 individuals) compared to other studies (max. genotype average = ∼3 individuals), which was likely due to the rather low abundance of aphids we observed (max. genotype average = ∼7 individuals).

Similar to arthropods, we found that direct environmental effects and host-plant genetic variation were the primary drivers of rhizosphere community assembly in the wind experiment. For example, the higher rarefied richness of rhizosphere bacteria on exposed plants appeared to be due to the higher concentrations of cations, such as Ca^2+^, Mg^2+^, and Cd^2+^, in the soil. These high concentrations of soil cations likely reflects a more alkaline soil pH, which is known to have a positive relationship with bacteria diversity (Fierer & Jackson 2006). The community composition of rhizosphere fungi, however, appeared more influenced by willow genotype. Although we failed to identify the heritable root traits mediating this interaction, we found that heritable variation in leaf flavonoid chemistry correlated with community responses. The composition of constituitive secondary metabolites is often highly correlated in leaf and root tissue (Kaplan *et al.* 2008), suggesting that plant secondary chemistry could influence the assembly of fungi on roots of different willow genotypes. Interestingly, leaf flavonoid chemistry was also important in mediating the assembly of arthropods in the ant-aphid experiment. An intriguing possibility is that leaf flavonoid chemistry plays an important role in the assembly of both above and belowground communities, but the direct effects of wind exposure on arthropods obscured this observation in the wind experiment. While measuring environmental effects on willow secondary metabolites was outside the scope of this study, our results suggest that secondary chemistry could be important in mediating plant genetic effects on above and belowground communities.

## Conclusion

Overall, our study reinforces the importance of host-plant genetic variation in shaping associated communities and extends this finding to natural biotic and abiotic gradients in coastal dunes. Our findings also suggest that predicting community responses to genetic and environmental variation is a complex task that may depend on historical processes that have shaped the genetic architecture for the populations of interest (i.e. sensitivity to specific environmental factors). Still, the effects of willow genetic variation were clear at both the level of plant traits and the community structure of foliar arthropods and fungi. Importantly, this suggests that host-plant evolution could have a strong influence on the biodiversity of above and belowground communities. Future studies should work toward understanding how these diverse communities feedback to impose selection pressures on host plants as well as how host-plant traits mediate interactions between above and belowground communities. In doing so, we will be able to work toward a more synthetic understanding of the evolutionary ecology of host plants and their diverse associated communities.

## Acknowledgments

We thank A. Pickart and the staff of Humboldt Bay National Wildlife Refuge (U.S. Fish and Wildlife Service) for permission to work at Lanphere Dunes and for facilitating experimental logistics. L. Mackas-Burns assisted with the fieldwork. We thank M.A.B.’s committee, M. O’Connor, D. Srivastava, and J. Brodie, whose helpful comments improved this manuscript. M.A.B. was supported by multiple fellowships from the University of British Columbia (BRITE Fellowship, James Robert Thompson Fellowship, and a Four-Year Fellowship). G.M.C. was supported by the Miller Institute for Basic Research in Science and a NSERC Discovery grant.

## Authors’ contributions

M.A.B., E.S.J., and G.M.C. conceived the ideas and designed the experiment; M.A.B. and G.M.C. designed and implemented the protocols for measuring the aboveground community, while S.E. and K.P. designed and implemented the protocols for measuring the belowground community; M.A.B., S.E., and B.L. collected the data; M.A.B. analyzed the data and led the writing of the manuscript. All authors contributed critically to the drafts and gave final approval for publication.

## Data accessibility

All data and R-scripts for reproducing the reported results are publicly available and can be found at the GitHub repository https://github.com/mabarbour/Lanphere_Experiments.git. The release associated with this version of the manuscript can be found at https://doi.org/10.5281/zenodo.840033. Data for these analyses will also be deposited on figshare.

## Online Appendix A: Supplementary Tables

**Table A1:**
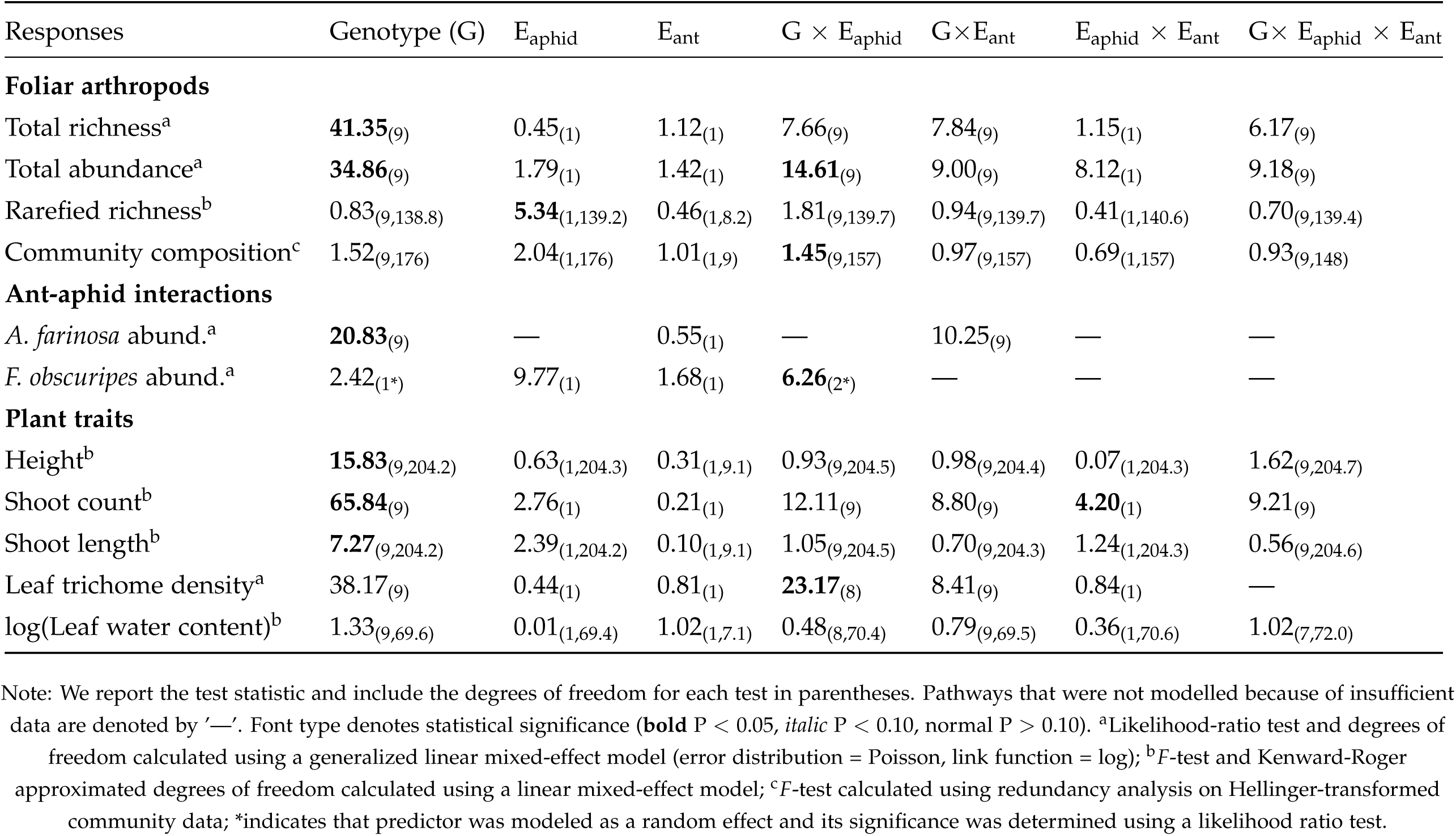
Summary of statistical models that analyze the effects of willow genotype, aphid treatment, and distance from ant mounds on the arthropod community, ant-aphid interactions, and plant traits.

| Responses | Genotype (G) | E<sub>aphid</sub> | E<sub>ant</sub> | G × E<sub>aphid</sub> | G × E<sub>ant</sub> | E<sub>aphid</sub> × E<sub>ant</sub> | G × E<sub>aphid</sub> × E<sub>ant</sub> |
| --- | --- | --- | --- | --- | --- | --- | --- |
| **Foliar arthropods** |  |  |  |  |  |  |  |
| Total richness<sup>a</sup> | **41.35**<sub>(9)</sub> | 0.45<sub>(1)</sub> | 1.12<sub>(1)</sub> | 7.66<sub>(9)</sub> | 7.84<sub>(9)</sub> | 1.15<sub>(1)</sub> | 6.17<sub>(9)</sub> |
| Total abundance<sup>a</sup> | **34.86**<sub>(9)</sub> | 1.79<sub>(1)</sub> | 1.42<sub>(1)</sub> | **14.61**<sub>(9)</sub> | 9.00<sub>(9)</sub> | 8.12<sub>(1)</sub> | 9.18<sub>(9)</sub> |
| Rarefied richness<sup>b</sup> | 0.83<sub>(9,138.8)</sub> | **5.34**<sub>(1,139.2)</sub> | 0.46<sub>(1,8.2)</sub> | 1.81<sub>(9,139.7)</sub> | 0.94<sub>(9,139.7)</sub> | 0.41<sub>(1,140.6)</sub> | 0.70<sub>(9,139.4)</sub> |
| Community composition<sup>c</sup> | 1.52<sub>(9,176)</sub> | 2.04<sub>(1,176)</sub> | 1.01<sub>(1,9)</sub> | **1.45**<sub>(9,157)</sub> | 0.97<sub>(9,157)</sub> | 0.69<sub>(1,157)</sub> | 0.93<sub>(9,148)</sub> |
| **Ant-aphid interactions** |  |  |  |  |  |  |  |
| *A. farinosa* abund.<sup>a</sup> | **20.83**<sub>(9)</sub> | — | 0.55<sub>(1)</sub> | — | 10.25<sub>(9)</sub> | — | — |
| *F. obscuripes* abund.<sup>a</sup> | 2.42<sub>(1*)</sub> | 9.77<sub>(1)</sub> | 1.68<sub>(1)</sub> | **6.26**<sub>(2*)</sub> | — | — | — |
| **Plant traits** |  |  |  |  |  |  |  |
| Height<sup>b</sup> | **15.83**<sub>(9,204.2)</sub> | 0.63<sub>(1,204.3)</sub> | 0.31<sub>(1,9.1)</sub> | 0.93<sub>(9,204.5)</sub> | 0.98<sub>(9,204.4)</sub> | 0.07<sub>(1,204.3)</sub> | 1.62<sub>(9,204.7)</sub> |
| Shoot count<sup>b</sup> | **65.84**<sub>(9)</sub> | 2.76<sub>(1)</sub> | 0.21<sub>(1)</sub> | 12.11<sub>(9)</sub> | 8.80<sub>(9)</sub> | **4.20**<sub>(1)</sub> | 9.21<sub>(9)</sub> |
| Shoot length<sup>b</sup> | **7.27**<sub>(9,204.2)</sub> | 2.39<sub>(1,204.2)</sub> | 0.10<sub>(1,9.1)</sub> | 1.05<sub>(9,204.5)</sub> | 0.70<sub>(9,204.3)</sub> | 1.24<sub>(1,204.3)</sub> | 0.56<sub>(9,204.6)</sub> |
| Leaf trichome density<sup>a</sup> | 38.17<sub>(9)</sub> | 0.44<sub>(1)</sub> | 0.81<sub>(1)</sub> | **23.17**<sub>(8)</sub> | 8.41<sub>(9)</sub> | 0.84<sub>(1)</sub> | — |
| log(Leaf water content)<sup>b</sup> | 1.33<sub>(9,69.6)</sub> | 0.01<sub>(1,69.4)</sub> | 1.02<sub>(1,7.1)</sub> | 0.48<sub>(8,70.4)</sub> | 0.79<sub>(9,69.5)</sub> | 0.36<sub>(1,70.6)</sub> | 1.02<sub>(7,72.0)</sub> |

Note: We report the test statistic and include the degrees of freedom for each test in parentheses. Pathways that were not modelled because of insufficient data are denoted by '—'. Font type denotes statistical significance (**bold** P < 0.05, *italic* P < 0.10, normal P > 0.10). <sup>a</sup>Likelihood-ratio test and degrees of freedom calculated using a generalized linear mixed-effect model (error distribution = Poisson, link function = log); <sup>b</sup>F-test and Kenward-Roger approximated degrees of freedom calculated using a linear mixed-effect model; <sup>c</sup>F-test calculated using redundancy analysis on Hellinger-transformed community data; \*indicates that predictor was modeled as a random effect and its significance was determined using a likelihood ratio test.

**Table A2:** Summary of statistical models that analyze the effects of willow genotype and wind exposure on soil characteristics and plant traits.

| Responses | Genotype (G) | E<sub>wind</sub> | E<sub>year</sub> | G × E<sub>wind</sub> | G × E<sub>year</sub> | E<sub>wind</sub> × E<sub>year</sub> | G × E<sub>wind</sub> × E<sub>year</sub> |
| --- | --- | --- | --- | --- | --- | --- | --- |
| **Soil characteristics** |  |  |  |  |  |  |  |
| Total N<sup>b</sup> | — | 5.08<sub>(1,9)</sub> | — | — | — | — | — |
| Soil moisture<sup>b</sup> | — | 3.52<sub>(1,9)</sub> | — | — | — | — | — |
| Percent organic matter<sup>b</sup> | — | 0.68<sub>(1,8.4)</sub> | — | — | — | — | — |
| Nutrient PC1<sup>b</sup> | — | 1.31<sub>(1,9)</sub> | — | — | — | — | — |
| **Plant traits** |  |  |  |  |  |  |  |
| Height<sup>b</sup> | **9.13**<sub>(9,145.3)</sub> | **29.10**<sub>(1,9.0)</sub> | 210.09<sub>(1,156.3)</sub> | 0.71<sub>(9,147.9)</sub> | 0.80<sub>(9,157.8)</sub> | **16.69**<sub>(1,158.4)</sub> | 1.84<sub>(9,160.9)</sub> |
| Shoot count<sup>a</sup> | **47.42**<sub>(9)</sub> | **9.91**<sub>(1)</sub> | 5.68<sub>(1)</sub> | 10.70<sub>(9)</sub> | **18.26**<sub>(9)</sub> | **12.53**<sub>(1)</sub> | 5.76<sub>(9)</sub> |
| Shoot length<sup>b</sup> | **4.97**<sub>(9,144.2)</sub> | **10.44**<sub>(1,9.0)</sub> | 75.36<sub>(1,158.5)</sub> | 0.84<sub>(9,146.9)</sub> | 1.61<sub>(9,160.1)</sub> | 0.05<sub>(1,160.7)</sub> | 0.70<sub>(9,163.2)</sub> |
| Leaf water content<sup>b</sup> | 4.90<sub>(9,129.0)</sub> | 0.97<sub>(1,8.7)</sub> | 2.93<sub>(1,139.7)</sub> | 0.47<sub>(9,132.0)</sub> | **2.80**<sub>(9,141.6)</sub> | 2.03<sub>(1,141.5)</sub> | 1.56<sub>(9,144.1)</sub> |
| Leaf trichome density<sup>b</sup><sub>2012</sub> | **67.31**<sub>(9)</sub> | 0.02<sub>(1)</sub> | — | 10.45<sub>(9)</sub> | — | — | — |
| SLA<sub>2013</sub><sup>b</sup> | **4.21**<sub>(9,122.5)</sub> | 0.34<sub>(1,8.9)</sub> | — | 1.19<sub>(9,123.4)</sub> | — | — | — |
| Leaf C:N<sub>2013</sub><sup>b</sup> | **4.88**<sub>(9,70.48)</sub> | 1.54<sub>(1,7.8)</sub> | — | 1.31<sub>(9,71.6)</sub> | — | — | — |
| Root C:N<sub>2013</sub><sup>b</sup> | 0.85<sub>(9,107.0)</sub> | 0.31<sub>(1,8.7)</sub> | — | 0.33<sub>(9,107.5)</sub> | — | — | — |

Note: We report the test statistic and include the degrees of freedom for each test in parentheses. Pathways that were not modelled because of insufficient data are denoted by '—'. Font type denotes statistical significance (**bold** P < 0.05, *italic* P < 0.10, normal P > 0.10). <sup>a</sup>Likelihood-ratio test and degrees of freedom calculated using a generalized linear mixed-effect model (error distribution = Poisson, link function = log); <sup>b</sup>F-test and Kenward-Roger approximated degrees of freedom calculated using a linear mixed-effect model.

**Table A3:**
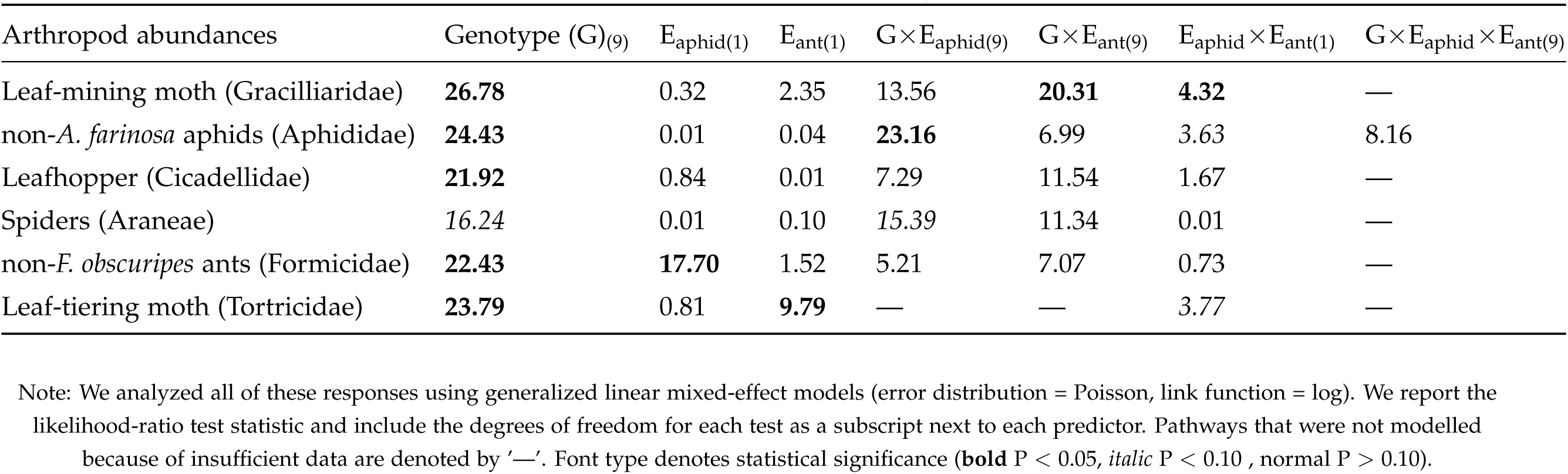
Summary of abundance responses of key arthropod guilds in the ant-aphid experiment.

**Table A4:** Summary of statistical models that analyze the effects of willow genotype and wind exposure on associated communities.

| Responses | Genotype (G) | E<sub>wind</sub> | E<sub>year</sub> | G × E<sub>wind</sub> | G × E<sub>year</sub> | E<sub>wind</sub> × E<sub>year</sub> | G × E<sub>wind</sub> × E<sub>year</sub> |
| --- | --- | --- | --- | --- | --- | --- | --- |
| **Foliar arthropods** |  |  |  |  |  |  |  |
| Richness<sup>a</sup> | **28.01**<sub>(9)</sub> | **10.33**<sub>(1)</sub> | **13.55**<sub>(1)</sub> | 3.74<sub>(9)</sub> | 9.85<sub>(9)</sub> | 0.92<sub>(1)</sub> | 7.04<sub>(9)</sub> |
| Abundance<sup>a</sup> | **25.25**<sub>(9)</sub> | **5.48**<sub>(1)</sub> | **6.72**<sub>(1)</sub> | 7.33<sub>(9)</sub> | 8.22<sub>(9)</sub> | 1.65<sub>(1)</sub> | 11.85<sub>(9)</sub> |
| Rarefied richness<sup>b</sup> | 1.96<sub>(9,71.1)</sub> | **22.82**<sub>(1,7.8)</sub> | 1.13<sub>(1,82.7)</sub> | 0.66<sub>(9,80.9)</sub> | — | 0.67<sub>(1,81.9)</sub> | — |
| Community composition<sub>2012</sub> | 0.96<sub>(9,51)</sub> | 1.26<sub>(1,7)</sub> | — | 0.91<sub>(9,42)</sub> | — | — | — |
| Community composition<sub>2013</sub> | 1.17<sub>(9,68)</sub> | **5.70**<sub>(1,9)</sub> | — | 0.69<sub>(6,62)</sub> | — | — | — |
| **Root-associated Mycorrhiza** |  |  |  |  |  |  |  |
| Rarefied richness<sub>2013</sub><sup>b</sup> | 0.87<sub>(9,95.1)</sub> | 0.88<sub>(1,8.8)</sub> | — | 0.93<sub>(9,95.9)</sub> | — | — | — |
| Community composition<sub>2013</sub> | **1.01**<sub>(9,117)</sub> | 1.18<sub>(1,9)</sub> | — | 0.87<sub>(9,108)</sub> | — | — | — |
| **Root-associated Bacteria** |  |  |  |  |  |  |  |
| Rarefied richness<sub>2013</sub><sup>b</sup> | 1.48<sub>(9,99.9)</sub> | **6.03**<sub>(1,7.8)</sub> | — | 1.35<sub>(9,100.5)</sub> | — | — | — |
| Community composition<sub>2013</sub> | 0.93<sub>(9,120)</sub> | 1.38<sub>(1,9)</sub> | — | 0.87<sub>(9,111)</sub> | — | — | — |

Note: We report the test statistic and include the degrees of freedom for each test in parentheses. Pathways that were not modelled because of insufficient data are denoted by '—'. Font type denotes statistical significance (**bold** P < 0.05, *italic* P < 0.10, normal P > 0.10). <sup>a</sup>Likelihood-ratio test and degrees of freedom calculated using a generalized linear mixed-effect model (error distribution = Poisson, link function = log); <sup>b</sup>F-test and Kenward-Roger approximated degrees of freedom calculated using a linear mixed-effect model; <sup>c</sup>F-test calculated using redundancy analysis on Hellinger-transformed community data.

**Table A5:** Summary of abundance responses of key arthropod guilds in the wind experiment.

| Arthropod abundances | Genotype (G)<sub>(9)</sub> | E<sub>wind(1)</sub> | G × E<sub>wind(9)</sub> | E<sub>year(1)</sub> | G × E<sub>year(9)</sub> | E<sub>wind</sub> × E<sub>year(1)</sub> | G × E<sub>wind</sub> × E<sub>year(9)</sub> |
| --- | --- | --- | --- | --- | --- | --- | --- |
| Leaf-mining moths (Gracillariidae) | **21.04** | **9.59** | — | 3.55 | — | 0.93 | — |
| Gall midges (Cecidomyiidae) | **24.89** | **19.49** | — | **238.13** | — | — | — |
| Leaf-tiering moths (Tortricidae) | **26.08** | 0.04 | 11.90 | **176.22** | — | 2.69 | — |
| Aphids<sub>2012</sub> (Aphididae) | 4.46 | 0.32 | — | — | — | — | — |
| Spiders (Araneae) | 9.36 | **6.58** | — | **9.79** | — | 0.20 | — |

Note: We analyzed all of these responses using generalized linear mixed-effect models (error distribution = Poisson, link function = log). We report the likelihood-ratio test statistic and include the degrees of freedom for each test as a subscript next to each predictor. Pathways that were not modelled because of insufficient data are denoted by '—'. Font type denotes statistical significance (**bold** P < 0.05, *italic* P < 0.10, normal P > 0.10).

**Table A6:** Summary of abundance responses of key fungal operational taxonomic units (OTUs) in the wind experiment.

| Fungal abundances | $E_{\text{wind}(1)}$ | Genotype (G)<sub>(9)</sub> | $G \times E_{\text{wind}(9)}$ |
| --- | --- | --- | --- |
| *Humicola nigrescens* (OTU 54) | 0.28 | **36.41** | — |
| *Ascomycota sp.* (OTU 68) | 0.18 | **24.38** | — |
| *Tremellomycetes sp.* (OTU 134) | **11.12** | **33.41** | — |
| *Cryptococcus terricola* (OTU 66) | 0.86 | **46.81** | **20.38** |
| *Cryptococcus albidus* (OTU 119) | 2.16 | **35.54** | 7.43 |
| *Cadophora finlandica* (OTU 262) | 0.29 | **43.37** | — |
| *Pseudogymnoascus roseus* (OTU 1606) | 0.01 | **22.70** | 13.58 |
| *Mortierella sp.* (OTU 113) | **4.14** | **21.08** | 9.21 |

Note: We analyzed all of these responses using generalized linear mixed-effect models (error distribution = Poisson, link function = log). We also specified an offset in the model (log(total fungal abundance)) to account for differences in sampling effort. We report the likelihood-ratio test statistic and include the degrees of freedom for each test as a subscript next to each predictor. Pathways that were not modelled because of insufficient data are denoted by '—'.
Font type denotes statistical significance (**bold** $P < 0.05$ , *italic* $P < 0.10$ , normal $P > 0.10$ ).

**Table A7:** Summary of loadings and variance explained by first two components from separate principal components analysis (PCA) of aboveground plant traits.

| Individual traits | Trait PC1 | Trait PC2 |
| --- | --- | --- |
| **Ant-aphid experiment** |  |  |
| Plant height | 0.51 | -0.49 |
| Shoot count | 0.43 | 0.25 |
| Shoot length | 0.64 | -0.14 |
| Leaf trichome density | -0.12 | -0.72 |
| Leaf water content | -0.36 | -0.39 |
| Variance explained | 39% | 29% |
| **Wind experiment, 2012** |  |  |
| Plant height | 0.55 | -0.18 |
| Shoot count | 0.47 | 0.29 |
| Shoot length | 0.68 | -0.07 |
| Leaf trichome density | -0.08 | 0.64 |
| Leaf water content | 0.09 | 0.69 |
| Variance explained | 36% | 24% |
| **Wind experiment, 2013** |  |  |
| Plant height | 0.46 | -0.29 |
| Shoot count | 0.49 | -0.27 |
| Shoot length | 0.45 | -0.20 |
| Leaf water content | -0.36 | -0.52 |
| Specific Leaf Area (SLA) | -0.46 | -0.42 |
| Leaf C:N | -0.04 | 0.59 |
| Variance explained | 45% | 26% |

Note: All traits were scaled to mean = 0 and SD = 1 prior to PCA to give them equal weight in the analysis.

**Table A8:** Summary of loadings and variance explained by first two components from principal components analysis (PCA) of soil properties.

| Soil properties | Soil PC1 | Soil PC2 |
| --- | --- | --- |
| $\text{NO}_3^-$ | -0.25 | -0.09 |
| $\text{NH}_4^+$ | -0.15 | -0.02 |
| $\text{Ca}^{2+}$ | 0.23 | 0.45 |
| $\text{Mg}^{2+}$ | 0.29 | 0.34 |
| $\text{K}^+$ | 0.23 | -0.20 |
| $\text{H}_2\text{PO}_4^-$ | 0.23 | 0.09 |
| $\text{Fe}^{3+}$ | 0.33 | -0.20 |
| $\text{Mn}^{2+}$ | 0.19 | 0.16 |
| $\text{Cu}^{2+}$ | 0.24 | -0.30 |
| $\text{Zn}^{2+}$ | 0.17 | -0.40 |
| $\text{B}(\text{OH})_4^-$ | 0.20 | -0.26 |
| $\text{SO}_4^{2-}$ | 0.26 | 0.10 |
| $\text{Pb}^{2+}$ | 0.21 | -0.10 |
| $\text{Al}^{3+}$ | 0.27 | -0.23 |
| $\text{Cd}^{2+}$ | 0.05 | 0.40 |
| Organic matter (%) | 0.35 | 0.14 |
| Soil moisture | 0.30 | 0.04 |
| Variance explained | 38% | 14% |

Note: All soil characteristics were scaled to mean = 0 and SD = 1 prior to PCA to give them equal weight in the analysis.

**Table A9:** Redundancy analyses of foliar arthropods and root-associated fungi and bacteria in the last year of the wind experiment.

| Community composition | $E_{\text{wind}}$ | Trait PC1 | Trait PC2 | Root C:N | Soil PC1 | Soil PC2 |
| --- | --- | --- | --- | --- | --- | --- |
| Arthropods | 0.83<sub>(1,9)</sub> | **12.05**<sub>(1,76)</sub> | 0.65<sub>(1,76)</sub> | — | — | — |
| Mycorrhiza | — | — | — | 1.17<sub>(1,115)</sub> | 1.89<sub>(1,8)</sub> | 0.85<sub>(1,8)</sub> |
| Bacteria | — | — | — | 1.31<sub>(1,116)</sub> | 1.90<sub>(1,8)</sub> | 0.81<sub>(1,8)</sub> |

Note: We report F-statistics and degrees of freedom in parenthesis. Effects that were not modelled because we did not hypothesize their direct effects are denoted by '—'. Font type denotes statistical significance (**bold** $P < 0.05$ , italic $P < 0.10$ , normal $P > 0.10$ ).

